# A single-domain response regulator activates exopolysaccharide synthesis by interacting with the initiating phosphoglycosyl transferase

**DOI:** 10.1101/2025.09.10.675344

**Authors:** Johannes Schwabe, Julia Monjaras-Feria, Timo Glatter, Patrick Blumenkamp, Oliver Rupp, Alexander Goesmann, Miguel A. Valvano, Lotte Søgaard-Andersen

## Abstract

Exported polysaccharides have crucial functions in bacteria. Polysaccharide biosynthesis in the ubiquitous Wzx/Wzy- and ABC-transporter-dependent pathways starts with the transfer of a sugar-1-phosphate from a nucleotide-sugar donor to undecaprenyl phosphate, a reaction catalyzed by a phosphoglycosyl transferase (PGT). Both reaction substrates are limited and shared among multiple glycoconjugate pathways, raising the question of how bacteria regulate these pathways. In *Myxococcus xanthu*s, EpsZ, which belongs to the family of large monotopic PGTs (monoPGTs), starts the Wzx/Wzy-dependent exopolysaccharide (EPS) biosynthesis. The Dif chemosensory system regulates EPS biosynthesis by an unknown mechanism *via* the phosphorylated single-domain response regulator EpsW (EpsW∼P). Here, we show that EpsW∼P stimulates EPS biosynthesis at the post-translational level. Moreover, MiniTurbo-based proximity labeling experiments suggest that EpsW∼P interacts directly with EpsZ. Additionally, heterologous expression of these two proteins in *Salmonella enterica* demonstrates that EpsW stimulates EpsZ enzymatic activity. *S. enterica* WbaP, the prototype large monoPGT, forms a functional homodimer, with dimerization involving a distinct cytoplasmic β-hairpin. However, AlphaFold-based structural modeling shows that EpsZ lacks this β-hairpin, suggesting an alternative mechanism for dimerization. Our structural modeling of a EpsZ-EpsW heterocomplex suggests that EpsW∼P, by direct interaction, promotes the formation of the stable EpsZ dimer. These findings uncover a new model for the regulation of polysaccharide biosynthesis in which EpsW∼P allosterically facilitates the formation of the active, dimeric conformation of EpsZ, thereby activating EPS biosynthesis at its initial step. Genomics and structural modeling suggest that the regulation of large monoPGTs by a single-domain response regulator is widespread in myxobacteria and potentially beyond.

**Significance:** Bacteria produce various polysaccharides with important biological functions and biotechnological applications. Polysaccharide synthesis is energy-costly and requires substrates that are in limited supply raising the question of how bacteria regulate these pathways. Here, we explored the regulation of exopolysaccharide biosynthesis in *Myxococcus xanthus*. We demonstrate that the phosphorylated single-domain response regulator EpsW activates exopolysaccharide biosynthesis at the post-translational level by stimulating the activity of the phosphoglycosyl transferase EpsZ. By directly interacting with EpsZ, phosphorylated EpsW facilitates the formation of the active, dimeric conformation of EpsZ, thereby activating exopolysaccharide biosynthesis at its initial step. We propose that this previously unrecognized regulatory mechanism is broadly conserved, not only in myxobacteria but also beyond.

## Introduction

Bacteria synthesize and export chemically diverse polysaccharides. These molecules contribute to biofilm formation, virulence, adhesion, motility, host-microbe interactions, and protection against phage infection and external stresses (1, 2), and also have applications in the food, biomedical, and pharmaceutical industries (3). Gram-negative bacteria synthesize and export these polysaccharides using three pathways: the Wzx/Wzy-, ABC transporter-, and synthase-dependent pathways (4–6). In the ubiquitous Wzx/Wzy- and ABC-transporter-dependent pathways, polysaccharide synthesis begins at the cytoplasmic leaflet of the inner membrane (IM) where an initiating phosphoglycosyl transferase (PGT) attaches a sugar-1-phosphate (sugar-1-P) from an activated nucleotide-sugar donor to undecaprenyl phosphate (Und-P) (7–9). The Und-PP-linked monosaccharide is the substrate for additional GTs that add monosaccharides and finalize the repeat unit in case of Wzx/Wzy-dependent pathways and the entire polysaccharide in case of ABC-transporter-dependent pathways (4). The monotopic PGTs (monoPGTs) constitute a prevalent PGT superfamily that shares a conserved catalytic domain (10, 11). MonoPGTs are further divided into small and large monoPGTs (9). Small monoPGTs, of which PglC from *Campylobacter concisus* (PglC*_Cc_*) is the prototype (11, 12), are monomeric and consist of only the catalytic domain (9, 13). By contrast, the large monoPGTs are dimeric and possess three conserved domains: a bundle of four transmembrane α-helices (TMHs), a cytoplasmic domain of unknown function (DUF), and the C-terminal catalytic domain (9, 14, 15). The *Salmonella enterica* WbaP (WbaP*_Se_*), which transfers galactose-1-phosphate (Gal-1-P) to Und-P, is the prototype of this family (16–20).

In the Wzx/Wzy-dependent pathways, the Wzx flippase translocates the Und-PP-linked repeat units across the IM into the periplasm, where the Wzy polymerase catalyzes their polymerization (4) by a process involving the polysaccharide co-polymerase (PCP) (4, 21). In pathways for capsular and secreted polysaccharides, the Und-PP-linked polysaccharides are exported across the outer membrane (OM) *via* a cell-envelope spanning export complex comprising either the PCP and an OM polysaccharide export (OPX) protein (4, 21) or a PCP, a periplasmic OPX protein, and an integral 16- to 18-stranded OM β-barrel protein (22, 23). Alternatively, in LPS biosynthetic pathways, the Und-PP-linked O-antigen polysaccharide is transferred to the lipid A-core oligosaccharide by the WaaL ligase (24) and transported to the cell surface by the Lpt lipopolysaccharide export pathway (25).

Polysaccharide biosynthesis *via* Wzx/Wzy-dependent pathways is energy-costly and also draws Und-P from a limited supply that must be sustained for the biosynthesis of other glycoconjugates, including the essential peptidoglycan (26–31). Accordingly, these pathways are regulated, commonly at the transcriptional level (32–38); but post-translational regulation has also been described, i.e. in *Staphylococcus aureus,* the bacterial tyrosine kinase (BYK) CapB1 phosphorylates the small monoPGT CapM, increasing its catalytic activity *in vitro* (39), and phosphoproteome analysis has revealed that the *Klebsiella pneumoniae* large monoPGT WcaJ is Tyr-phosphorylated albeit a functional analysis of this modification is lacking (40). While the post-translational mechanism(s) of regulation of Wzx/Wzy-dependent pathways are incompletely understood, the post-translational regulation of synthase-dependent pathways is well characterized, frequently involving the second messenger c-di-GMP (6). In these cases, c-di-GMP can stimulate synthase activity by binding to an allosteric site (41), to a non-catalytic partner protein of the synthase (42–44), or by enabling the interaction of the catalytic domain and a non-catalytic partner protein (45).

In the Gram-negative bacterium *Myxococcus xanthus*, the exopolysaccharide (EPS), which is synthesized and exported by the Wzx/Wzy-dependent EPS pathway (22, 23, 46–49), is crucial for type IV pili (T4P)-dependent motility, adhesion, development, and biofilm formation (50). EPS synthesis starts by the transfer of Gal-1-P to Und-P, catalyzed by the large monoPGT EpsZ (46) (Figure 1A). EPS biosynthesis is regulated by the Dif chemosensory system (51–53) *via* the phosphorylated single-domain response regulator EpsW (54) (Figure 1A). The DifE histidine protein kinase phosphorylates EpsW (54), but how phosphorylated EpsW (EpsW∼P) stimulates EPS biosynthesis is unknown. DifE also phosphorylates the single-domain response regulator DifD, which, together with the DifG phosphatase, acts as a phosphate sink competing with EpsW for phosphorylation by DifE to inhibit EPS biosynthesis (55, 56) (Figure 1A). The signal(s) regulating Dif activity are unknown, but it has been suggested that T4P extension activates the Dif system (57–59).

**Figure 1.**
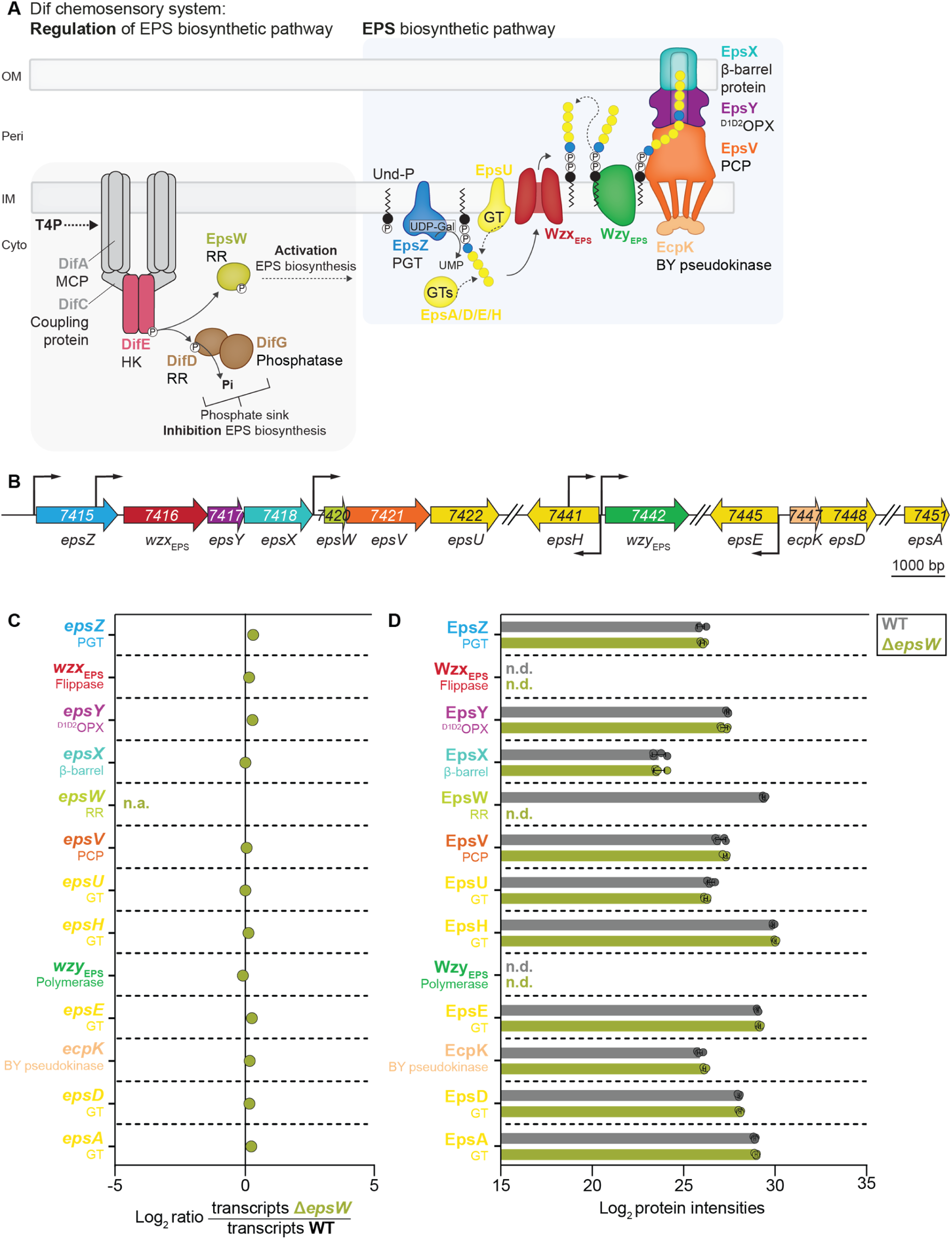
The Eps proteins accumulate independently of EpsW. (A) Schematic of the Dif chemosensory system and the EPS biosynthetic pathway in *M. xanthus*. Stippled line, mechanism unknown; continuous line, mechanism known. See text for details. MCP, methyl-accepting chemotaxis protein; HK, histidine kinase; RR, response regulator. EpsZ transfers Gal-1-P to Und-P. The repeat units are built by several glycosyl transferases (GT) and translocated into the periplasm by the Wzx_EPS_ flippase. The Wzy_EPS_ polymerase builds the final EPS molecule. EpsV is the polysaccharide co-polymerase (PCP) and functions together with the bacterial tyrosine pseudokinase (BY pseudokinase) EcpK (49). EPS is exported across the OM through a bipartite translocon comprising the ^D1D2^OPX protein EpsY, an OM polysaccharide export protein only comprising the periplasmic domains D1 and D2, and the 18-stranded OM β-barrel protein EpsX (22, 23). (B) Operon structure of the two gene clusters for *eps* biosynthesis. Kinked arrows, transcription start sites (95). Numbers, MXAN_locus tags. EpsW, while a downstream target of the Dif chemosensory system, is encoded in the *eps* locus. (C) Differential expression of *eps* genes in the Δ*epsW* mutant compared to WT. The RNA-seq experiment was performed with four biological replicates per strain from cells grown in suspension culture. X-axis, log_2_-fold ratio of the mean transcripts in the Δ*epsW* mutant over the mean transcripts in the WT calculated using the DESeq2 method (96). n.a., non-applicable. Statistical analysis was performed in the DeSeq2 analysis. No significant differences were identified (adjusted *P-*value ≤ 0.001). (D) Protein abundance in whole-cell proteomes of the Δ*epsW* mutant compared to WT. The LFQ-MS-based proteomics was performed with four biological replicates per strain from cells grown as in (C). X-axis, normalized log_2_ intensities of Eps proteins. Data points represent each of the four biological replicates. Error bars, standard deviation (SD). No significant differences were identified (Welch’s test, *P-*value ≤ 0.001). n.d., not detected.

Here, we elucidate how EpsW∼P activates EPS biosynthesis. Using global transcriptomic and proteomic analyses, we demonstrate that EpsW regulates EPS biosynthesis at the post-translational level. MiniTurbo-based proximity labeling experiments in *M. xanthus* strongly suggest that EpsW∼P directly interacts with EpsZ. Additionally, *in vivo* heterologous expression experiments in *S. enterica* demonstrated that while EpsZ alone had low Gal-1-P transferase activity, this activity was strongly stimulated when co-expressed with EpsW, suggesting that EpsW∼P, by direct interaction, brings about the active EpsZ conformation. Computational structural analyses using AlphaFold indicated that while EpsZ alone may be compromised in dimer formation, EpsW, by directly interacting with EpsZ, promotes the formation of the stable EpsZ dimer. Together, these findings support a previously undescribed allosteric mechanism whereby a single-domain response regulator, by direct interaction, stimulates the enzymatic activity of a large monoPGT by facilitating its active, dimeric conformation.

## Results

### EpsW activates EPS biosynthesis at the post-translational level

Single-domain response regulators can function in phosphorelays to regulate gene expression or engage in protein-protein interactions to control the activity of a downstream protein (complex) (60, 61). To explore how EpsW stimulates EPS biosynthesis, we identified differentially expressed genes and differentially accumulating proteins by RNA sequencing (RNA-seq) and whole-cell, label-free quantitative (LFQ) mass spectrometry-based proteomics in the Δ*epsW* mutant and the parental wild-type (WT) strain. In these experiments, cells were grown exponentially in suspension, a condition where *M. xanthus* synthesizes and exports EPS (46, 62).

We first analyzed genes and proteins known to be involved in EPS biosynthesis or its regulation, including the core genes/proteins for T4P formation and function. RNA transcript levels of all *eps* genes (Figure 1B and C), *dif* genes (Figure S1A), other genes encoding regulators of EPS biosynthesis (Figure S1B), and genes for the core proteins for T4P formation and function (Figure S2A) were similar in WT and the Δ*epsW* mutant. The corresponding proteins detected by whole-cell proteomics accumulated at similar or slightly higher levels in the Δ*epsW* mutant compared to WT (Figure 1D, Fig. S1C and D; Figure S2B). These findings suggest that EpsW does not activate EPS biosynthesis by regulating the abundance of Eps and Dif proteins, other regulators of EPS biosynthesis, or the core proteins for T4P formation and function.

T4P extension requires a priming complex made of four minor pilins and the PilY1 adhesin (63). The *M. xanthus* genome contains three gene clusters encoding distinct sets of minor pilins/PilY1 proteins (63). Cluster_1 alone and cluster_3 alone are sufficient for T4P formation, while cluster_2 does not contribute to T4P formation, and the corresponding proteins do not detectably accumulate (63). RNA transcripts of all *cluster_1*, *cluster_2,* and *cluster_3* genes were detected in the WT (Figure S3A), with low levels of *cluster_2*, as exemplified by *pilY1.2* (Figure S3B). In the Δ*epsW* mutant, the transcript levels of *cluster_1* genes were as in WT, except for *pilX1,* which was slightly but significantly reduced (Figure S3A). The transcript levels of all five *cluster_3* genes and two *cluster_2* genes were also significantly reduced in the Δ*epsW* mutant (Figure S3A). The transcript levels of *hsfA* and *hsfB*, encoding a phosphorelay that regulates transcription of *cluster_1* and *cluster_3* expression (64), were slightly but significantly increased in the Δ*epsW* mutant (Figure S3A). However, whole-cell proteomics revealed significantly reduced abundance of only two cluster_3 proteins, while the abundance of HsfA was slightly increased (Figure S3C). To assess whether these changes in the abundance of cluster_3 minor pilins could cause the EPS biosynthetic defect in the Δ*epsW* mutant, we determined EPS biosynthesis in different cluster mutants (Figure S3D). While the Δ*cluster_2*Δ*cluster_3* double mutant synthesized EPS at WT levels, the Δ*cluster_1*Δ*cluster_2*Δ*cluster_3* triple mutant, which lacks all genes encoding the three priming complexes, did not synthesize EPS (Figure S3D). Thus, cluster_1 is sufficient for WT levels of EPS biosynthesis, arguing that the EPS biosynthesis defect in the Δ*epsW* mutant is not due to the reduced accumulation of two of the cluster_3 proteins.

We also investigated changes at the global transcript and protein levels in the Δ*epsW* mutant compared to the WT (Materials and Methods). Except for the *cluster_3* genes (Figure S3A), only two genes encoding hypothetical proteins had reduced transcript levels (Table S1). Four genes, encoding CRISPR-associated proteins, had increased transcript levels (Table S1). A total of 22 and five proteins had reduced and increased accumulation levels, respectively (Table S2). However, none of these genes and proteins have been implicated in EPS biosynthesis.

Collectively, these findings suggest that the EPS biosynthetic defect in the absence of EpsW does not depend on altered gene expression or differences in protein abundance, indicating that EpsW stimulates EPS biosynthesis at the post-translational level.

### Phosphorylation of EpsW enhances the EpsW-EpsZ interaction *in vivo*

We speculated that EpsW could stimulate EPS biosynthesis post-translationally by engaging in direct protein-protein interaction(s). Therefore, we searched for EpsW interaction partners *in vivo* by proximity labeling using EpsW fused to the biotin ligase miniTurbo (mTurbo) (65–68). We ensured that mTurbo-based proximity labeling could detect Eps proteins and DifE (the only known interaction partner of EpsW) by assessing the presence of surface-accessible cytoplasmic Lys residues in these proteins. Manual counting using AlphaFold2-generated models revealed that all Eps proteins and DifE, except for EpsY and EpsX, which lack cytoplasmic domains (Figure 1A) (22, 23), have Lys residues amenable to biotinylation (Table S3).

For the biotinylation experiments, we ectopically expressed *mTurbo-epsW-FLAG* from the *pilA* promoter in three different mutants: Δ*epsW*, Δ*difE*Δ*epsW* (lacking the DifE kinase that phosphorylates EpsW) and Δ*difD*Δ*difG*Δ*epsW* (lacking DifD and the DifG phosphatase that jointly function as a phosphate sink for the phosphoryl-groups of DifE) (Figure 1A). As controls for unspecific biotinylation, we co-expressed sfGFP-mTurbo-FLAG from the vanillate-inducible promoter (65) and untagged EpsW from the *pilA* promoter in the three mutants. mTurbo-EpsW-FLAG and sfGFP-mTurbo-FLAG (in the presence of 7.5 µM vanillate) accumulated at similar levels, and both had biotin-ligase activity in all three strains (Figure S4A−D). Moreover, mTurbo-EpsW-FLAG restored EPS biosynthesis and T4P-dependent motility in the Δ*epsW* and the Δ*difD*Δ*difG*Δ*epsW* mutants, with the Δ*difD*Δ*difG*Δ*epsW* strain producing slightly more EPS (Figure S5), demonstrating that the fusion protein is functional and there is enhanced phosphate flux from DifE to mTurbo-EpsW-FLAG in the absence of DifD/DifG. As expected, mTurbo-EpsW-FLAG supported neither EPS biosynthesis nor T4P-dependent motility in the Δ*difE*Δ*epsW* strain (Figure S5). Similarly, the control strains co-expressing untagged EpsW and sfGFP-mTurbo-FLAG in the Δ*epsW* and the Δ*difD*Δ*difG*Δ*epsW* strains had EPS biosynthesis and T4P-dependent motility, again with slightly more EPS biosynthesis in the Δ*difD*Δ*difG*Δ*epsW* strain (Figure S5). As expected, the Δ*difE*Δ*epsW* strain co-expressing untagged EpsW and sfGFP-mTurbo-FLAG did not have EPS biosynthesis and T4P-dependent motility (Figure S5).

In the proximity labeling experiments in the WT background (Figure 2A, Table S4), 11 proteins, including EpsW, were significantly enriched in the mTurbo-EpsW-FLAG samples, indicating successful biotinylation and enrichment of the fusion protein. Importantly, the DifE kinase, the only known EpsW interaction partner, was enriched. Strikingly, EpsZ was also enriched. None of the remaining eight proteins (Table S4), including the glycosyl hydrolase (family 57) MXAN_2994 and the GNAT-family acetyltransferase MXAN_4576 that may participate in monosaccharide synthesis or modification, have been identified as important for EPS biosynthesis.

**Figure 2.**
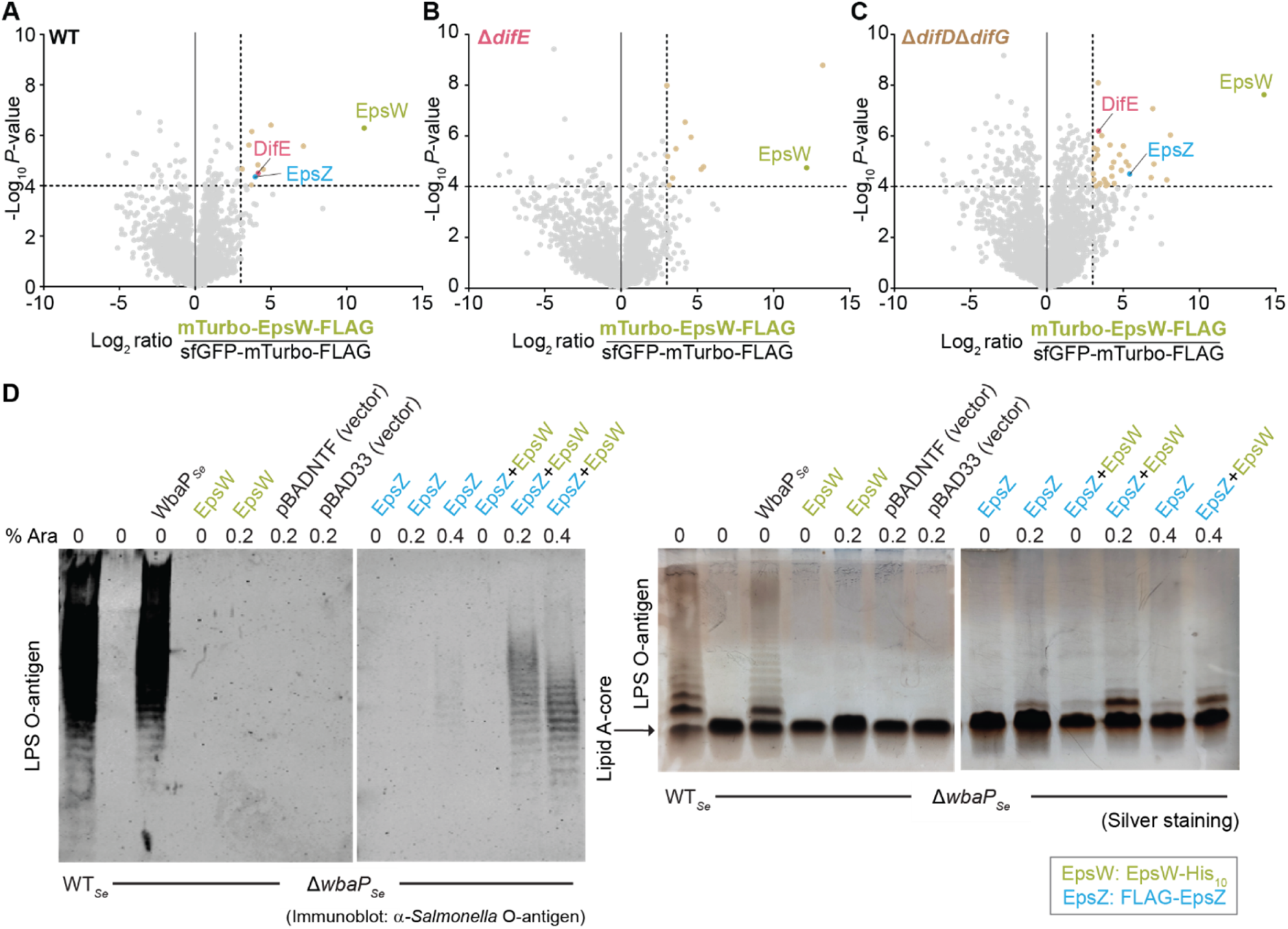
EpsW phosphorylation enhances the EpsW-EpsZ interaction *in vivo* and EpsW stimulates EpsZ Gal-1-P transferase activity. (A-C) Volcano plots showing the enrichment of proteins in the proximity labeling experiment with mTurbo-EpsW-FLAG and sfGFP-TurboID-FLAG variants in the otherwise WT (A), Δ*difE* (B), and Δ*difD*Δ*difG* (C) strains. *mTurbo-epsW-FLAG* was expressed from the *pilA* promoter on a plasmid integrated in a single copy at the Mx8 *attB* site. In the *sfGFP-mTurbo-FLAG* strains, *epsW* was expressed from the *pilA* promoter on a plasmid integrated in a single copy at the Mx8 *attB* site, and *sfGFP-mTurbo-FLAG* was expressed under control of the vanillate-inducible promoter from a plasmid integrated in a single copy at the *18-19* intergenic locus. For all strains, samples were prepared from four biological replicates. X-axis, log_2_-fold ratios of mean protein intensities in mTurbo-EpsW-FLAG samples relative to the sfGFP-mTurbo-FLAG samples. Y-axis, −log_10_ *P*-value. Significantly enriched proteins fulfill the criteria log_2_ fold ratio ≥3.0; −log_10_ *P*-value ≥4.0, and are indicated by wheat-colored data points and listed in Table S4 to Table S6. Significance thresholds are indicated with dashed lines. Data points represent the means of four biological replicates. (D) EpsW stimulates EpsZ Gal-1-P transferase activity. Complementation of the *S. enterica* Δ*wbaP* mutant (Δ*wbaP_Se_*) by ectopic expression of the indicated proteins. LPS samples were extracted from the indicated strains and examined by immunoblotting using α-*Salmonella* O-antigen antiserum (left) and silver staining (right). Samples from an equal amount of cells were loaded per sample. Arabinose (Ara) was added as indicated. FLAG-EpsZ and EpsW-His_10_ were expressed from pMP146 (vector: pBADNTF) and pJSc143 (vector pBAD33), respectively under the control of an arabinose-inducible promoter. WbaP*_Se_* was expressed from pSM13 under the control of a constitutive *lac* promoter.

Eleven proteins including EpsW, but not EpsZ were significantly enriched (Figure 2B, Table S5) in the Δ*difE* mutant (no EpsW phosphorylation). None of the remaining ten proteins have been identified as important for EPS biosynthesis (Table S5).

In the absence of DifD and DifG (increased EpsW phosphorylation), 33 proteins, including EpsW, DifE and EpsZ were significantly enriched (Figure 2C, Table S6), with EpsZ enrichment even more prominent (log_2_-fold increase of 1.4 compared to EpsZ enrichment in the WT strain). None of the remaining proteins, including the GNAT-family acetyltransferase MXAN_2367, have been identified as important for EPS biosynthesis (Table S6).

Among the four proteins that were enriched in both WT and Δ*difD*Δ*difG* strains but not in the Δ*difE* strain (Table S4, S5, and S6), only DifE and EpsZ were reported to function in EPS biosynthesis, and the two remaining proteins are not predicted to function in polysaccharide biosynthesis. Because the labeling radius of mTurbo *in vivo* is ∼10 nm (69), these results indicate that upon phosphorylation by DifE, EpsW is in close proximity to and interacts with EpsZ. Because an EpsW variant with a substitution of the phosphorylatable Asp residue (D^58^) does not stimulate EPS biosynthesis (54), these observations imply that EpsW∼P stimulates EPS biosynthesis by interacting with EpsZ.

### EpsW activates EpsZ Gal-1-P transferase activity in a heterologous system

We used a *S. enterica*-based heterologous expression system to test whether EpsW stimulates EpsZ Gal-1-P transferase activity *in vivo*. In *S. enterica*, WbaP*_Se_* initiates the synthesis of the LPS O-antigen by catalyzing the transfer of Gal-1-P to Und-P (19). Using LPS O-antigen biosynthesis as a readout, we previously showed that EpsZ with an N-terminal FLAG-tag (FLAG-EpsZ) weakly complements the *S. enterica* Δ*wbaP* (Δ*wbaP_Se_*) mutant, but these earlier experiments did not include EpsW (46).

Here, we expressed FLAG-EpsZ without or with an active EpsW-His_10_ variant (Figure S6A and B) in the Δ*wbaP_Se_* mutant. As a positive control, we initially confirmed that the ectopic expression of WbaP restored LPS O-antigen synthesis in this mutant (Figure 2D). As previously observed, FLAG-EpsZ alone, expressed from an arabinose-inducible promoter, only weakly complemented the LPS O-antigen biosynthetic defect in the presence of 0.4% arabinose, as shown in immunoblots with *Salmonella* O-antigen-specific antibodies and by silver staining (46) (Figure 2D). As expected, EpsW-His_10_, which was also expressed from an arabinose-inducible promoter, did not restore LPS O-antigen synthesis (Figure 2D). By contrast, co-expression of FLAG-EpsZ and EpsW-His_10_ in the presence of 0.2% and 0.4% arabinose efficiently restored LPS O-antigen biosynthesis in the Δ*wbaP_Se_* mutant (Figure 2D). Immunoblots using α-FLAG antibodies demonstrated that FLAG-EpsZ abundance correlated with the arabinose concentration and was independent of the presence or absence of EpsW-His_10_ (Figure S6C). Under the conditions used for protein denaturation and SDS-PAGE, FLAG-EpsZ accumulated predominantly in the monomeric form and less as higher molecular weight oligomers in both strains, while other large monoPGTs display a more equal distribution between the monomer and the oligomers (20, 70, 71). In immunoblots, we were unable to detect EpsW-His_10_ in the Δ*wbaP_Se_* mutant using α-His_6_ antibodies.

We conclude from these results that EpsW stimulates EpsZ Gal-1-P transferase activity *in vivo*. In *M. xanthus*, EpsW-dependent stimulation of EPS biosynthesis depends on the DifE-dependent phosphorylation (54). Therefore, we speculate that EpsW in the Δ*wbaP_Se_* mutant is either non-specifically phosphorylated by a *S. enterica* histidine protein kinase or autophosphorylates using low molecular weight phosphodonor(s) like acetyl phosphate or carbamoyl phosphate (72, 73). Because EpsW∼P is in close proximity to and likely interacts with EpsZ in *M. xanthus*, these observations suggest that EpsW∼P enhances EpsZ Gal-1-P transferase activity by direct interaction.

### EpsZ has a similar catalytic domain as prototypic PGTs but lacks a characteristic β-hairpin in the cytoplasmic DUF

Our results argue that EpsW∼P is a direct activator of EpsZ enzymatic activity. However, the activity of the PglC*_Cc_* and WbaP*_Se_* prototype PGTs does not depend on regulatory proteins, raising the question of the EpsZ-specific mechanism of activation. To understand the mechanism of EpsW∼P-mediated EpsZ activation, we performed structural modeling in three steps.

First, we generated AlphaFold2 models of monomeric WbaP*_Se_* and EpsZ. The WbaP*_Se_* model had high confidence (Figure S7A and B), revealing a structure consisting of the catalytic domain, TMH bundle, and cytoplasmic DUF (Figure 3A), as previously described (14, 20). The EpsZ model, also of high confidence (Figure S7A and C), contained equivalent domains (Figure 3A). Next, we compared the solved structure of monomeric PglC*_Cc_* (11), which only consists of the conserved catalytic domain, with the high-confidence structural models of WbaP*_Se_* and EpsZ (Figure 3A).

**Figure 3.**
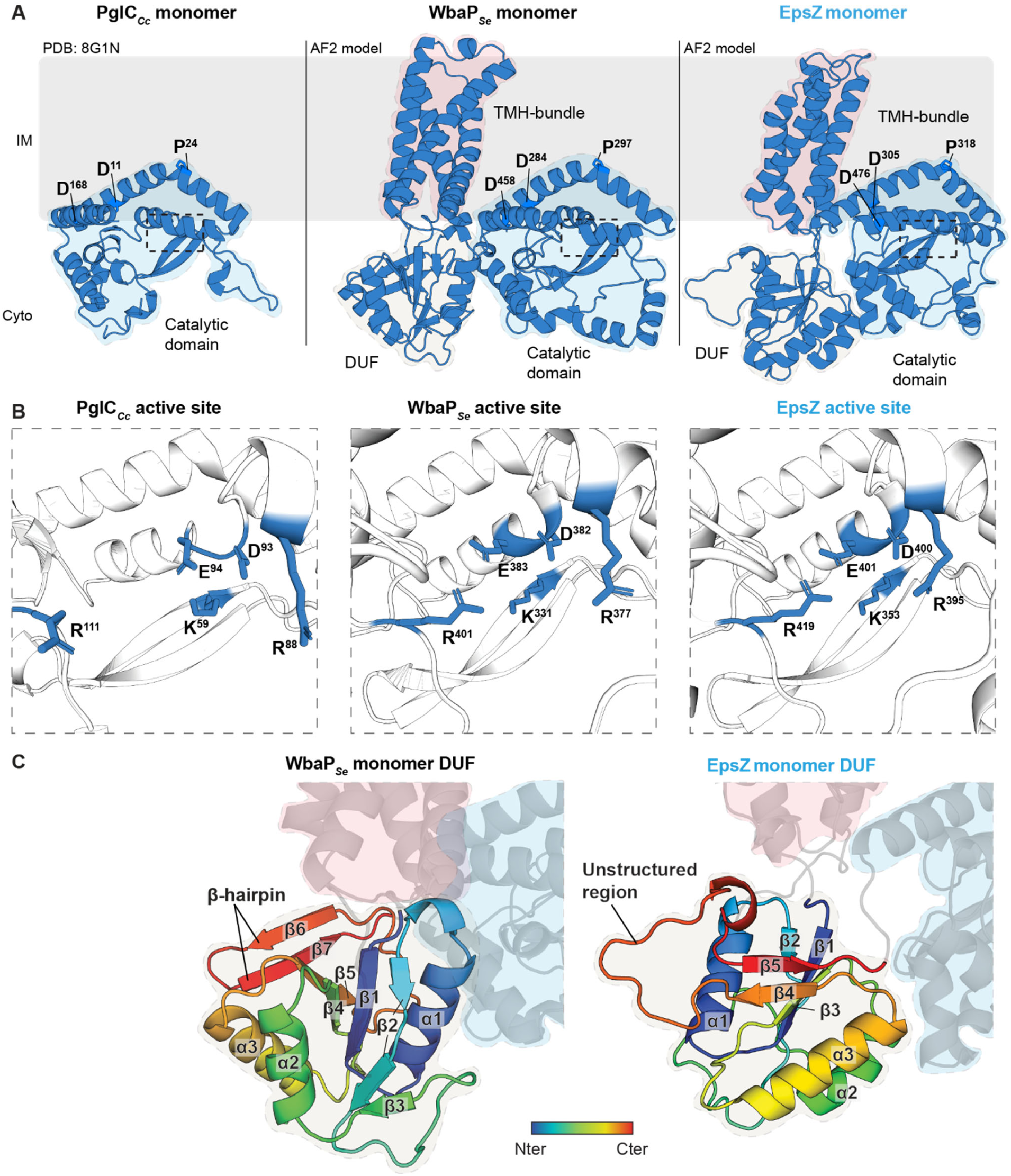
EpsZ is a large monoPGT lacking a characteristic β-hairpin in the cytoplasmic DUF. (A) Computational structural comparison of EpsZ with the prototype monoPGTs PglC*_Cc_* and WbaP*_Se_*. Left panel, solved structure of the small monoPGT PglC*_Cc_* (11). Middle and right panels, rank 1 AlphaFold2 models of monomeric WbaP*_Se_* and monomeric EpsZ. All panels, regions in light blue indicate catalytic domain. Regions in silk and light red in the middle and right panels indicate the cytoplasmic DUF and the TMH bundle. Conserved residues important for the re-entrant α-helix architecture are highlighted, see text for details. The stippled boxes mark the position of the active site. Protein orientation in the IM is based on computational analysis using the PPM server (97). (B) Computational structural comparison of the active sites of EpsZ with those of PglC*_Cc_* and WbaP*_Se_.* Left panel, active site of the solved structure of PglC*_Cc_* (11). Middle and right panels, active sites of the rank 1 AlphaFold2 models of monomeric WbaP*_Se_* and monomeric EpsZ. Conserved residues important for catalytic activity are highlighted (11, 13, 16). (C) Computational structural comparison of the cytoplasmic DUF of EpsZ with that of WbaP*_Se_*. Left and right panels, cytoplasmic DUF of the rank 1 AlphaFold2 models of monomeric WbaP*_Se_* and EpsZ. In both panels, the DUF is shown using a gradient from blue (N-terminus) to red (C-terminus). Numbering of α-helices and β-sheets is included. In both panels, regions colored in blue and red indicate the catalytic domain and the TMH bundle.

In monoPGTs, the catalytic domain integrates into the IM *via* the re-entrant α-helix, which is kinked by a Pro residue and enters and exits the IM’s lipid bilayer at the cytoplasmic leaflet (10, 11). This Pro residue is part of the D_X12_P motif, which is strictly conserved in monoPGTs (74), and also present in PglC*_Cc_*, WbaP*_Se,_* and EpsZ (10, 11, 46). In the WbaP*_Se_* and EpsZ structural models, the D_X12_P motif mapped to similar positions (D^284^ and P^297^ in WbaP*_Se_*, D^305^ and P^318^ in EpsZ) on the re-entrant α-helix as in the catalytic domain of PglC*_Cc_* (D^11^ and P^24^) (Figure 3A). Additionally, a strictly conserved Asp residue suggested to function in stabilizing the re-entrant α-helix in PglC*_Cc_* (D^168^) (11, 74) mapped to equivalent positions in WbaP*_Se_* (D^458^) and EpsZ (D^476^) (Figure 3A).

In PglC*_Cc_*, the transfer of the sugar-1-P to Und-P at the active site involves a catalytic dyad (D^93^E^94^) and additional residues for substrate orientation (K^59^, R^88^, R^111^) (11) that are conserved among monoPGTs (74) (Figure 3B). The WbaP*_Se_* and EpsZ structural models had active site architectures comparable to that of PglC*_Cc_* (Figure 3B). This agrees with the current model that the catalytic mechanism of monoPGTs is conserved irrespective of the nucleotide-sugar substrates (74–76). Therefore, we conclude that the activation of EpsZ is not related to differences in its catalytic domain, which is similar to the homologous domains in PglC*_Cc_* and WbaP*_Se_*.

The previous results suggested that the EpsZ-specific mechanism for activation involves other EpsZ domains. Therefore, we compared the cytoplasmic DUF of EpsZ and WbaP*_Se_*. The WbaP*_Se_* cytoplasmic DUF model shows a central β-sheet composed of five β-strands (β1-β5), with β2 interrupted by a short unstructured region (Figure 3C). Three α-helices surround the central β-sheet (Figure 3C). Additionally, the C-terminal region of the DUF contains a β-hairpin built by the β6/β7-strands (Figure 3C). Despite the cytoplasmic DUF in EpsZ having a superficially similar fold to its WbaP*_Se_* counterpart, with both featuring a central β-sheet built of five β-strands surrounded by three α-helices, (Figure 3C), three key differences exist: (i) unlike WbaP*_Se_*, the EpsZ cytoplasmic DUF lacks the C-terminal β-hairpin (Figure 3C), (ii) the β4/β5-strands are separated by an unstructured region (Figure 3C), and (iii) the α2/α3-helices are oriented towards the catalytic domain, whereas in WbaP*_Se_*, it is the α1-helix that faces the catalytic domain.

### EpsZ alone might be compromised in dimer formation

In the second step of the structural modeling, we considered that recent structural characterization and computational modeling suggest that WbaP*_Se_* forms a homodimer (14, 15), stabilized by the two cytoplasmic DUFs *via* the C-terminal β-hairpins built from the β6/β7-strands (Figure 3C) (14). The absence of this DUF β-hairpin in the high-confidence model of the EpsZ monomer, as indicated above (Figure 3C), suggests that EpsZ dimer formation could be compromised.

The structure of the WbaP*_Se_* homodimer has been solved at ∼4.1 Å resolution (14). Therefore, we generated and compared AlphaFold2-Multimer models of WbaP*_Se_* and EpsZ dimers. WbaP*_Se_* was modeled as a homodimer (Figure 4A) with high confidence and low predicted alignment error (pAE) (Figure S8A and B). The predicted interface template modeling score (ipTM) of 0.79 is close to the stringent 0.80 cutoff (77), supporting an overall accurate prediction of the dimer interface (Figure 4B). PDBePISA (78) analysis calculated a dimer interface area of 4098 Å^2^ (Figure 4B). As reported earlier (14), we observed that the two cytoplasmic DUFs contribute significantly to the dimer interface, with the β-hairpin of one protomer’s DUF participating in β-sheet augmentation (79) with the other protomer’s β-sheet and *vice versa* (Figure 4C). Moreover, the tip of one protomer’s β-hairpin is close to the other protomer’s catalytic domain and *vice versa* (Figure 4C). Additionally, the two protomers interface in a region involving the TMH bundles (Figure 4B).

**Figure 4.**
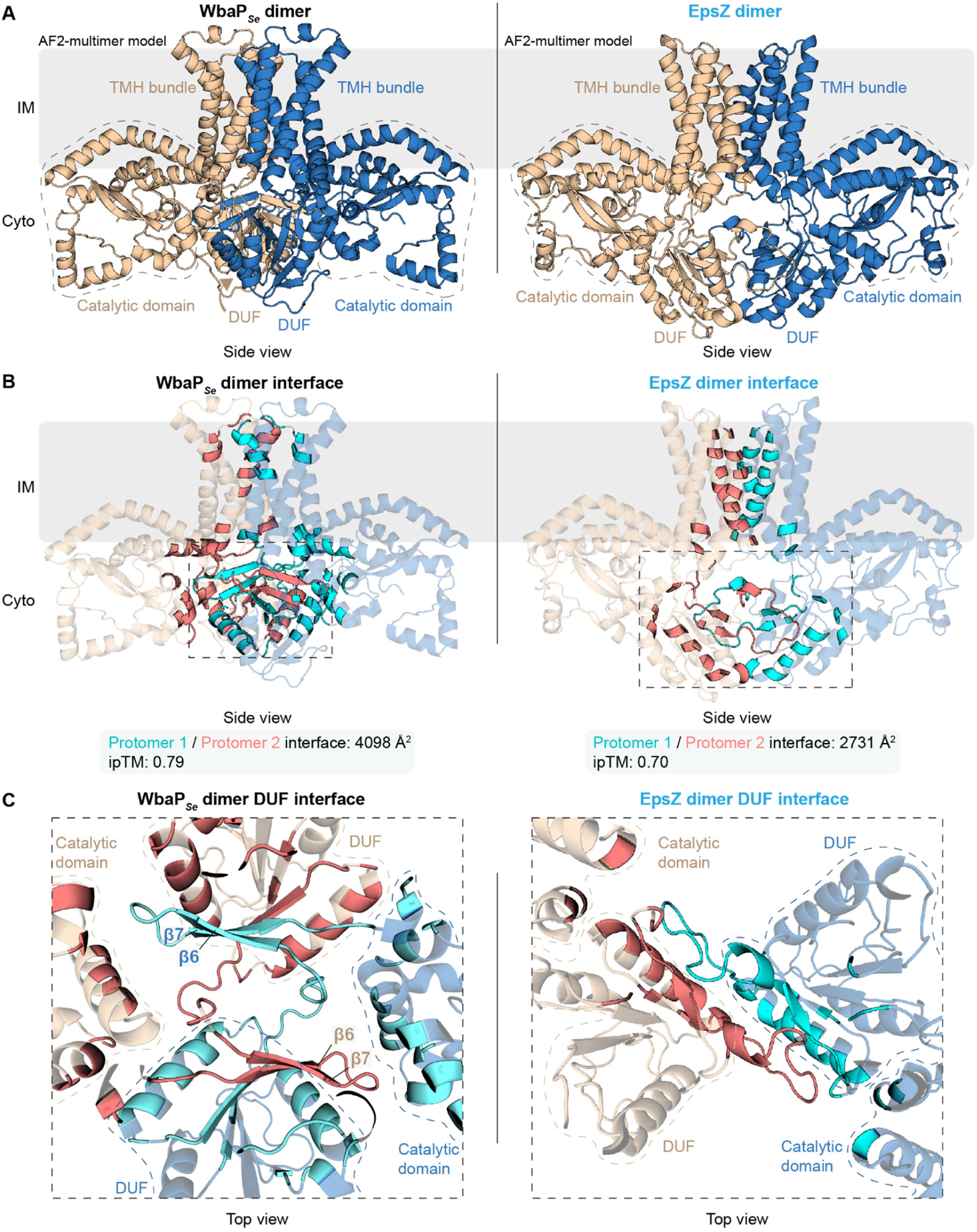
EpsZ alone may be compromised in dimer formation. (A) Computational structural comparison of WbaP*_Se_* and EpsZ dimers. Left and right panels, rank 1 AlphaFold2-Multimer models of dimeric WbaP*_Se_* and dimeric EpsZ. Individual protomers are colored blue and wheat. Regions with stippled outlines indicate the catalytic domains. The cytoplasmic DUFs and the TMH bundles are marked. The position of the proteins in the IM was computed using the PPM server (97). (B) Analysis of the dimer interface of the rank 1 AlphaFold2-Multimer models of the WbaP*_Se_* and EpsZ dimers. Both panels, residues at the interface between protomers are highlighted in cyan (contributed by the blue-colored protomer) and salmon (contributed by the wheat-colored protomer). Below, the overall surface area of the interface between the protomers and the corresponding ipTM value are shown. Both panels, the stippled area marks the interface within the cytoplasmic DUFs of the two protomers. The interface analysis was performed *via* PDBePISA (78). (C) Top view of zoomed-in image of the interface between the cytoplasmic DUFs of the protomers in the WbaP*_Se_* and EpsZ dimers. The zoomed region is marked by a stippled box in (B). Both panels, residues at the dimer interface are highlighted as described in (A), and stippled lines indicate the catalytic domains and DUFs of each protomer. Left panel, the characteristic β6/β7-hairpins involved in β-sheet augmentation are marked.

The model of the EpsZ homodimer also displayed a dimeric arrangement with the TMH bundles and the cytoplasmic DUFs in proximity (Figure 4A); however, the PDBePISA analysis calculated a reduced dimer interface (2731 Å^2^) compared to the WbaP*_Se_* dimer (4098 Å^2^) (Figure 4B), and no β-sheet augmentation in the DUFs’ interface (Figure 4C). Moreover, the central part of the dimer, particularly the interface between the two DUFs, was only modeled with moderate confidence based on pLDDT (Figure S8A and C). Additionally, the pAE for the EpsZ dimer was higher compared to the WbaP*_Se_* dimer, indicating reduced confidence in the model’s spatial accuracy (Figure S8C, compare to Figure S8B). Finally, the ipTM value for the EpsZ dimer was 0.70, indicating a less confident interface prediction than for the WbaP*_Se_* dimer. Together, we conclude that the absence of the β-hairpin and the inability to model the EpsZ dimer with high confidence indicate that EpsZ alone cannot form a stable dimer.

### EpsW may facilitate the active, dimeric conformation of EpsZ

Our experimental data indicates that EpsZ interacts with EpsW∼P. Therefore, in the third step of the structural modeling, we explored an interaction between EpsZ and EpsW. To this end, we first generated a model of EpsW using AlphaFold2. Notably, it is not possible to model post-translational modifications (e.g., phosphorylation of D^58^ in EpsW that is phosphorylated by DifE (54)) using AlphaFold2 (80). Additionally, introducing substitutions in the sequence used for model generation (i.e., that could mimic phosphorylation) only has a limited impact on models compared to using the native sequences (81, 82). Thus, while EpsW∼P is required to activate EPS biosynthesis in *M. xanthus* (54) and is the form of EpsW that interacts with EpsZ based on the proximity labeling experiments (Figure 2A-C), we used the native EpsW sequence for modeling. The high-confidence EpsW model (Figure S8D and E) showed that the EpsW monomer adopts the characteristic structure of the receiver domain of response regulators (83), consisting of a central five-stranded parallel β-sheet surrounded by five α-helices (Figure 5A). A structure homology-based search using Foldseek (84) identified the BeF ^-^-activated form of the single-domain response regulator CheY, which mimics the phosphorylated form of CheY*_E. coli_*, in complex with CheZ (PDB: 1KMI (85)), with high confidence as the closest structural homolog of EpsW in the PDB^100^ database (Probability 1.0, sequence identity 29.4%, e-value 7.14^-12^). This suggests that our EpsW model represents the activated phosphorylated state of the molecule.

**Figure 5.**
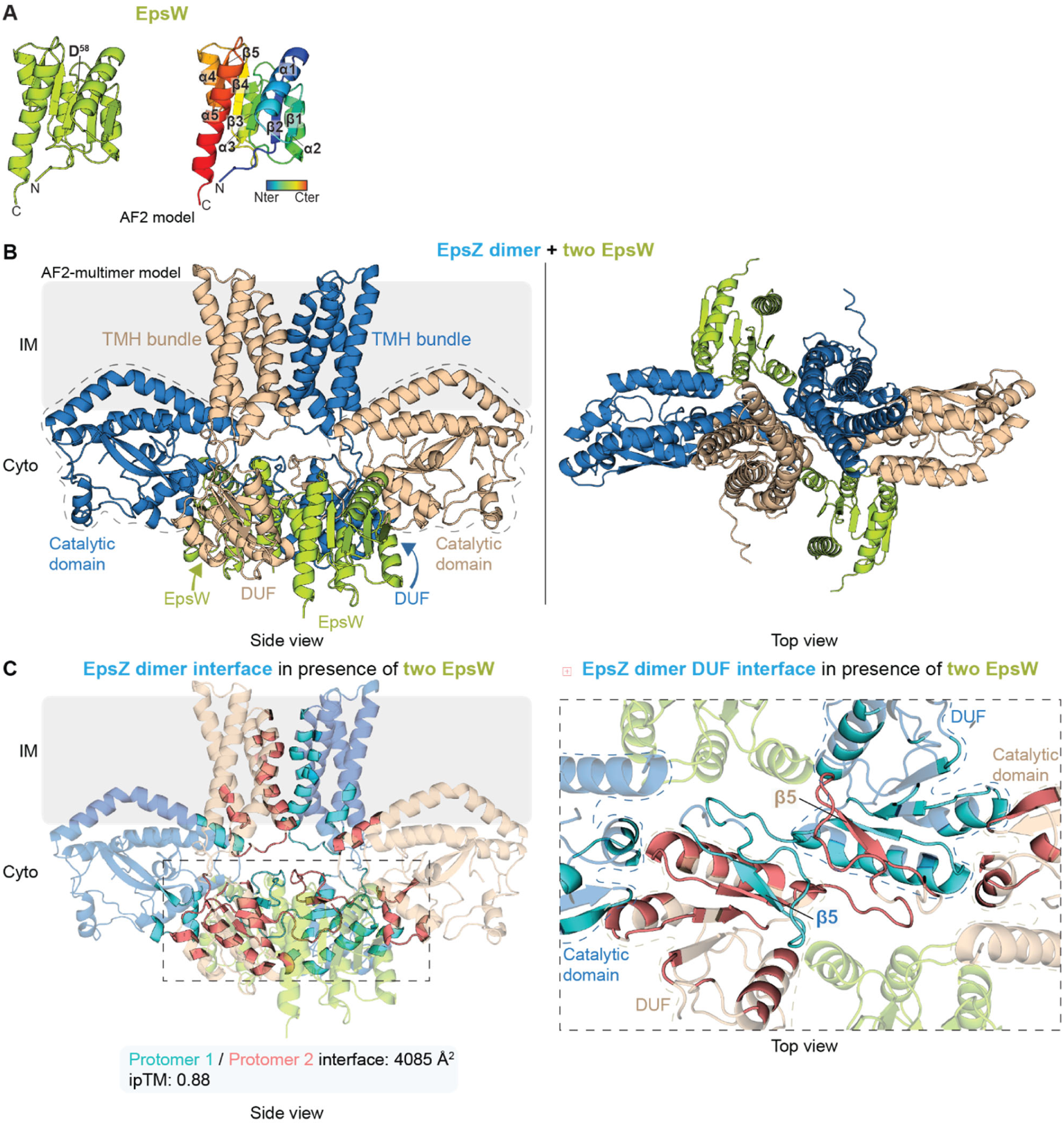
EpsW may facilitate the active, dimeric conformation of EpsZ. (A) Rank 1 AlphaFold2 model of EpsW. Left panel, EpsW in lime-green. The phosphorylated D^58^ based on (54) is indicated. Right panel, protein depicted in a gradient from blue (N-terminus) to red (C-terminus). Numbering of α-helices and β-strands is included. (B) Computational AlphaFold2-Multimer structural model of the EpsW-EpsZ heterocomplex in 2:2 stoichiometry. EpsW colored lime-green and EpsZ protomers in blue and wheat. Model rank 1 is shown. Regions with stippled outlines indicate the catalytic domains; the cytoplasmic DUFs and the TMH bundles are marked. (C) Analysis of the EpsZ dimer interface within the AlphaFold2-Multimer model of the EpsW-EpsZ heterocomplex. Left panel, residues at the interface between the EpsZ protomers are highlighted in cyan (contributed by the blue-colored protomer) and salmon (contributed by the wheat-colored protomer). Below, the overall surface area of the interface between the EpsZ protomers and the corresponding ipTM value is shown. Right panel, zoomed-in top view of the interface between the cytoplasmic DUFs of the EpsZ protomers in the EpsW-EpsZ heterocomplex. The residues at the dimer interface are highlighted as in the left panel, and stippled lines indicate the catalytic domains and DUFs of each protomer. EpsW is colored lime-green. The β-sheet augmentation of the β5-strands is marked.

Next, we generated a model comprising two copies of EpsZ and two copies of EpsW using AlphaFold2-Multimer. In the EpsZ-EpsW heterocomplex, the two EpsW copies were modeled symmetrically on either side of the cytoplasmic part of the EpsZ dimer (Figure 5B). The EpsZ-EpsW model was of high confidence, and with a low pAE and high ipTM score of 0.88 (Figure S8F and G; Figure 5C), indicating a highly confident prediction of the interfaces within the heterocomplex (77). The modeled structure of EpsW alone is similar to that of EpsW in complex with EpsZ (RMSD: 0.251, C_α_ 831), suggesting that EpsW in this model is also in its activated, phosphorylated state.

Finally, we investigated the predicted interface between the EpsZ protomers in the EpsZ-EpsW heterocomplex. The EpsZ-EpsZ interface was located between the two DUFs and the TMH bundles (Figure 5C). The catalytic domains of the EpsZ protomers were swapped relative to the TMH bundles in the EpsZ-EpsW heterocomplex (Figure 5B and C) when compared to the EpsZ dimer without EpsW (Figure 4A and B). This resulted in an increased interface area between the EpsZ protomers to 4085 Å^2^ (Figure 5C). Moreover, in the presence of two EpsW copies, we observed β-sheet augmentation within the EpsZ-EpsZ interface between the two DUFs (Figure 5D). Specifically, the β5-strand of one protomer’s DUF (Figure 3C) folds into the β-sheet of the adjacent protomer’s DUF and *vice versa* (Figure 5C). We note that this interaction is different from that in the WbaP*_Se_* dimer, where the DUFs’ β-hairpins mediate β-sheet augmentation, rather than a strand of the central β-sheet (Figure 4C). Importantly, EpsW does not contribute to the active site in EpsZ.

Together, our results from structural modeling predict a heterocomplex of two copies each of EpsZ and EpsW, in which EpsW directly interacts with EpsZ. Because EpsW is likely modeled in its phosphorylated state and does not contribute to the EpsZ active site, we conclude that EpsW∼P is a direct allosteric activator of EpsZ that facilitates the dimeric, active conformation of EpsZ, thereby stimulating Gal-1-P transferase activity and initiation of EPS biosynthesis.

### The EpsZ-EpsW mechanism may be conserved in orthologous Wzx/Wzy-dependent pathways

To investigate whether the EpsZ-EpsW mechanism is widespread, we searched for orthologous Wzx/Wzy-dependent EPS biosynthesis pathways in the 21 fully sequenced myxobacterial genomes (Table S7). We identified 15 EpsZ orthologs (23, 46) of which nine co-occurred with an EpsW ortholog (Figure 6A). Next, we performed structural modeling in the same three steps as for EpsZ/EpsW.

**Figure 6.**
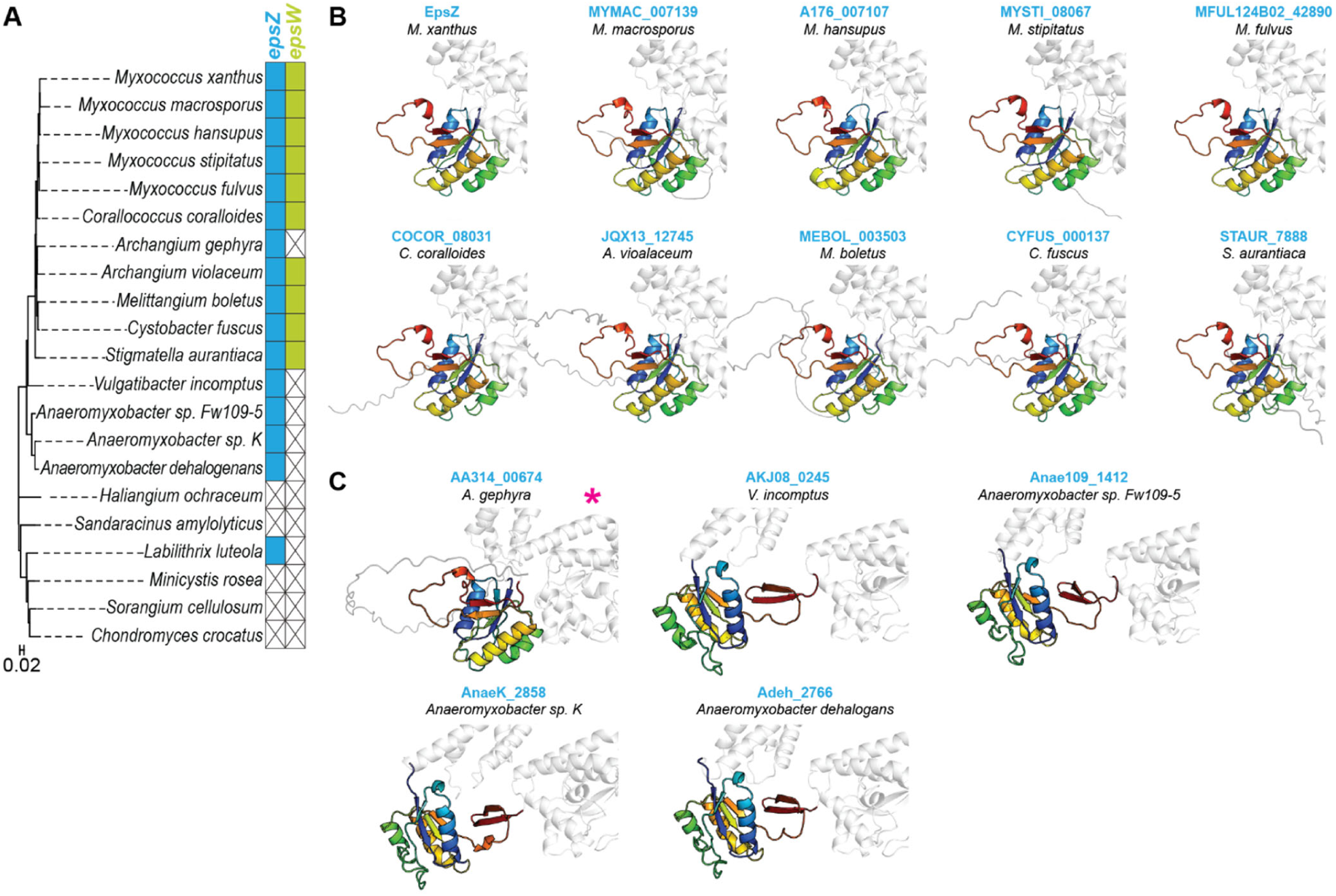
The absence of the β-hairpin in myxobacterial EpsZ orthologs largely correlates with the presence of an EpsW ortholog. (A) Left panel, phylogenetic tree of myxobacteria with fully-sequenced genomes based on 16S rRNA sequences. Right panels, co-occurrence of EpsZ and EpsZ orthologs (blue) and EpsW and EpsW orthologs (lime-green) in myxobacteria as identified in (23, 46). The absence of orthologs is indicated by an X. (B) Computational structural comparison of the cytoplasmic DUFs of EpsZ and the nine myxobacterial EpsZ orthologs co-occurring with an EpsW ortholog. In all panels, the cytoplasmic DUF of the rank 1 AlphaFold2 models of the full-length proteins is highlighted using a gradient from blue (N-terminus) to red (C-terminus). (C) Computational structural comparison of the cytoplasmic DUFs of myxobacterial EpsZ orthologs not co-occurring with an EpsW ortholog. In all panels, the cytoplasmic DUF of the rank 1 AlphaFold2 model is shown. The DUF is shown as in (B). * indicates that the *A. gephyra* EpsZ ortholog does not have a β-hairpin.

The models of the monomeric structures of the EpsZ orthologs were, generally, of high confidence (Figure S9 and S10A). Only the *Labilitrix luteola* EpsZ ortholog was modeled with low confidence and has a DUF with an unusual structure comprising additional α-helices and β-strands (Figure S10B); therefore, this protein was excluded in further analyses. Comparing the cytoplasmic DUFs of the remaining protein models, we found that this domain lacked the β-hairpin in ten of the EpsZ orthologs and instead had an unstructured region connecting two of the β-sheet’s strands (Figure 6B and C). The β-hairpin was present in four EpsZ orthologs (Figure 6C). The EpsZ orthologs without the β-hairpin co-occurred with EpsW orthologs, except for *Archangium gephyra*, whereas EpsZ orthologs with the β-hairpin did not co-occur with EpsW orthologs (Figure 6A-C). Thus, the absence or presence of the β-hairpin in these EpsZ orthologs almost perfectly correlates with the presence or absence of an EpsW ortholog.

Dimeric models of the nine EpsZ orthologs that co-occurred with an EpsW ortholog (Figure 7A) were overall of mediocre confidence (Figure S11). In five of the nine dimers, the domain arrangement resembled that of the EpsZ dimer, but in the remaining four models, the catalytic domains were swapped relative to the TMH bundles (Figure 7A). Finally, we modeled the nine heterocomplexes containing two copies of the EpsZ ortholog and two copies of the corresponding EpsW ortholog. In these nine models, the two copies of the EpsW orthologs were positioned symmetrically on either side of the cytoplasmic part of the dimer of the EpsZ orthologs, reminiscent of the heterocomplex in *M. xanthus* (Figure 7B). Except for the *Melittangium boletus* model, the models of the heterocomplexes had high ipTM scores and low pAE values (Figure 7C and S12), indicating high confidence in the predicted interactions in the heterocomplexes. Moreover, the ipTM scores and pAE values of the heterocomplexes were markedly higher than those of the corresponding homodimeric complexes (Figure 7C, Figure S11 and S12).

**Figure 7.**
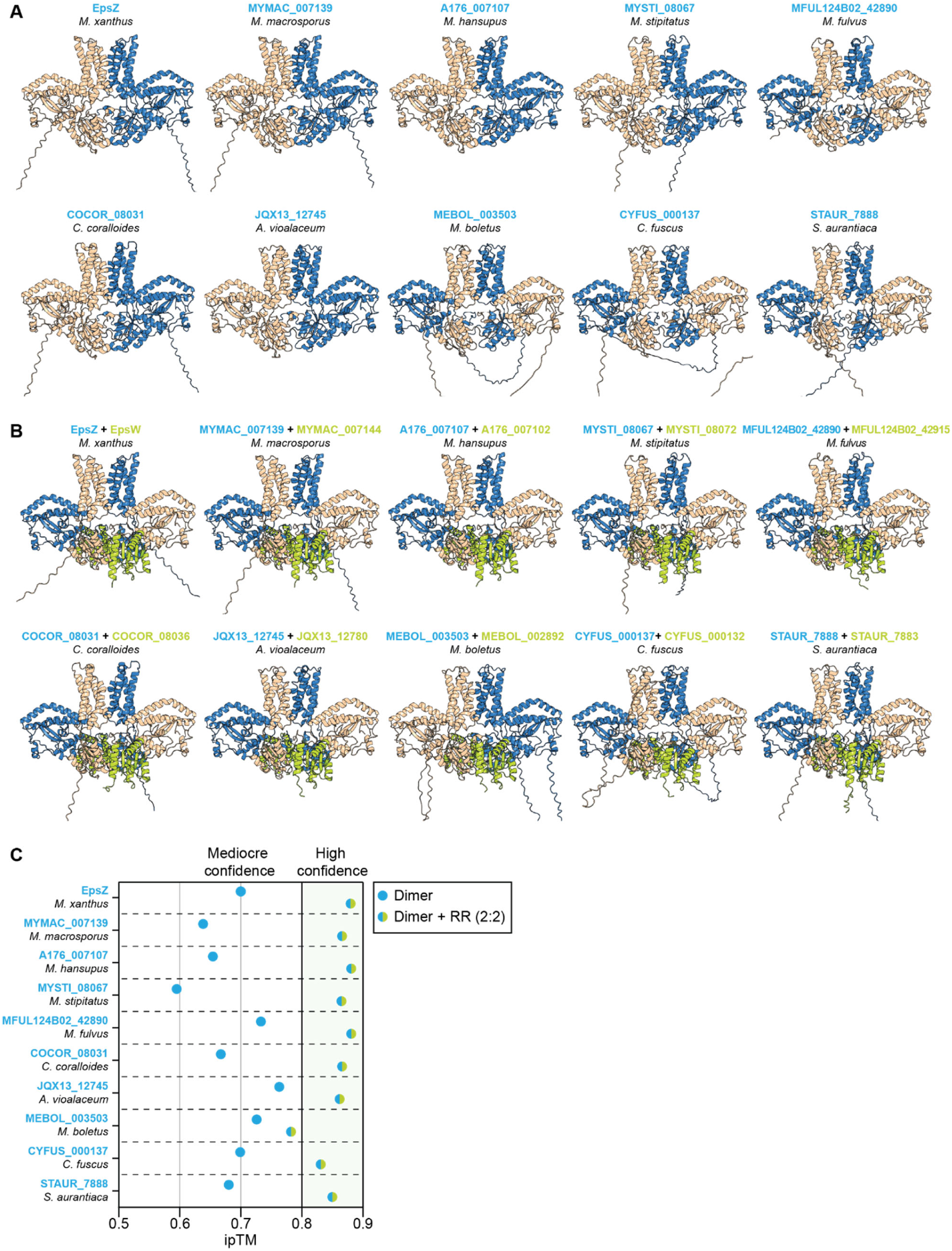
The EpsW/EpsZ mechanism may be conserved in myxobacteria. (A) Computational structural comparison of dimers of EpsZ and the nine myxobacterial EpsZ orthologs co-occurring with an EpsW ortholog. In all panels, rank 1 AlphaFold2-Multimer model of the dimeric protein is shown, and the individual protomers are colored blue and wheat. (B) Computational structural comparison of dimers EpsZ and nine myxobacterial EpsZ orthologs together with the corresponding EpsW ortholog in 2:2 stoichiometry. In all panels, rank 1 AlphaFold2-Multimer model of the heterocomplex is shown, the individual EpsZ ortholog protomers are colored blue and wheat, and the two copies of the EpsW ortholog in lime-green. (C) ipTM values of the rank 1 AlphaFold2-Multimer models in (A) and (B). X-axis, data points represent the ipTM values of the rank 1 model of the dimer of EpsZ and the EpsZ orthologs, either without (points colored blue) or with the EpsW ortholog (points colored blue/lime-green). An ipTM score above 0.80 indicates a confident interface prediction, scores between 0.60 and 0.80 fall into a grey zone where predictions may or may not be correct, and scores below 0.60 indicate a failed prediction (77).

Collectively, these computational analyses suggest that the direct interaction of a single-domain response regulator (i.e., EpsW and its orthologs) and a large monoPGT lacking the characteristic β-hairpin (i.e., EpsZ and its orthologs) is conserved in several other myxobacteria, suggesting that the EpsW/EpsZ mechanism for activating a large monoPGT is widespread.

## Discussion

This study elucidated how the single-domain response regulator EpsW, upon phosphorylation by DifE, activates EPS biosynthesis. Three key observations support the conclusion that EpsW∼P activates EPS biosynthesis by stimulating EpsZ Gal-1-P transferase activity. First, global transcriptomic and proteomic analyses indicate that EpsW∼P stimulates EPS biosynthesis at the post-translational level. Second, mTurbo-based proximity labeling in *M. xanthus* reveals that EpsW∼P is in close proximity to and likely directly interacts with EpsZ. Third, *in vivo* heterologous expression experiments in *S. enterica* show that the low level of Gal-1-P transferase activity of EpsZ alone is significantly enhanced by co-expressing EpsW.

Computational structural biology analyses suggest that EpsZ, similar to other large monoPGTs, has the C-terminal canonical catalytic domain partially embedded in the cytoplasmic leaflet of the IM, an N-terminal TMH bundle, and a central cytoplasmic DUF. However, the cytoplasmic DUF of EpsZ lacks the characteristic β-hairpin that contributes most of the dimer interface in the homodimeric WbaP*_Se_* and stabilizes the dimer by β-sheet augmentation (14). This difference was reflected in the lower confidence of the EpsZ homodimer model, and the smaller interface area compared to dimeric WbaP*_Se_*. By contrast, the model of the EpsZ-EpsW heterocomplex with 2:2 stoichiometry was of high confidence. In this complex, EpsZ had a dimeric arrangement, and each of the two EpsW copies symmetrically and directly contacts an EpsZ protomer’s DUF and catalytic domain on both sides of the EpsZ dimer. In the EpsZ-EpsW heterocomplex, the two catalytic domains of EpsZ were swapped relative to the two TMH bundles. Moreover, the EpsZ dimer interface area in the heterocomplex configuration increased to a similar level as in the WbaP*_Se_* dimer. Based on these models, we posit that EpsZ alone cannot efficiently dimerize and that EpsW∼P is an allosteric regulator, which facilitates the formation of the dimeric conformation of EpsZ upon direct interaction. Because EpsW∼P activates EpsZ Gal-1-P transferase activity, we suggest that this dimeric conformation, analogous to the functional WbaP*_Se_* homodimer, represents the active state of EpsZ. Similar conclusions can be drawn for myxobacterial EpsZ orthologs and their partner EpsW orthologs since structural modeling also supports with high confidence the formation of heterocomplexes reminiscent of the *M. xanthus* EpsZ/EpsW heterocomplex. Together, these findings support that a phosphorylated single-domain response regulator stimulates, by direct protein-protein interaction, the activity of a large monoPGT in several myxobacteria.

Post-translational modification of the *S. aureus* small monoPGT CapM by Tyr phosphorylation was suggested to stimulate enzymatic activity (39). Our results demonstrate a novel mode of post-translational regulation in which a regulatory protein stimulates the initiating large monoPGT *via* direct protein-protein interaction. The EpsZ-EpsW mechanism also adds to the list of mechanisms for post-translational regulation of the initial step of polysaccharide biosynthesis as described for synthase-dependent pathways (41–45). Because accumulation of the Eps proteins is independent of EpsW, we speculate that the EpsW-dependent mechanism provides *M. xanthus* with the ability to swiftly switch on the EPS pathway in response to T4P extension and other stimuli potentially sensed by the Dif chemosensory system. We also note that EPS biosynthesis is energy-costly and uses the lipid carrier Und-P, which is required for the biosynthesis of the essential peptidoglycan and other polysaccharides in *M. xanthus* (50). Thus, it is likely beneficial to regulate EPS biosynthesis at the initial step rather than in further downstream steps, in which repeat units would accumulate without being incorporated into the final EPS chain, and Und-P would be titrated away from other biosynthetic pathways.

Based on structural modeling, we suggest that stimulation of a large monoPGT by a phosphorylated single-domain response regulator domain is likely conserved across several myxobacteria, supporting that it could be more widespread in bacteria. Interestingly, a recent sequence-based bioinformatic analysis revealed complexity in the domain architecture of the small monoPGT family (13). These monoPGTs may incorporate additional domains, such as sugar-modifying and regulatory domains, including receiver domains of response regulators (13). However, because small monoPGTs are functional monomers (11), the regulation mechanism may not involve dimer formation. Yet, these observations indicate complex regulatory and functional diversity in the initial step of polysaccharide biosynthesis, supporting the idea that receiver domains could be involved in regulating monoPGTs in other organisms.

## Supporting information

All supplementary information

## Acknowledgment

We thank María Perez-Burgos and Dorota Skotnicka for plasmids and *M. xanthus* strains, Dobromir Szadkowski for providing a Matlab script to help with illustrating the AlphaFold model confidence plots, and Luca Blöcher for initial experiments. The Max Planck Society supported this work.

## Conflict of Interest

The authors declare no conflict of interest.

## Data Availability

The data supporting the findings of this study are all included in the manuscript and its supplementary file. The count table and mapping results of the RNA-seq experiment have been deposited at EBI ArrayExpress under accession number E-MTAB-14794. The mass spectrometry data of whole cell proteomics and proximity labeling experiments have been deposited to the ProteomeXchange Consortium *via* the PRIDE partner repository with the dataset identifiers PXD063981 and PXD063973, respectively. All materials are available from the corresponding author.

## Materials and Methods

### Strains and cell growth

All *M. xanthus* strains used in this study are derivatives of the WT strain DK1622 (86) and are listed in Table S7. Plasmids and oligonucleotides are listed in Table S8 and Table S9, respectively. In-frame deletions in *M. xanthus* were constructed *via* two-step homologous recombination as described (87). Plasmids for complementation experiments were integrated in a single copy by site-specific recombination at the Mx8 *attB* site or by homologous recombination at the *18-19* intergenic locus. All plasmids were verified by DNA sequencing, and all strains were verified by PCR. *M. xanthus* cultures were grown at 32°C in 1% CTT broth (1% [w/vol] Bacto casitone, 10 mM Tris-HCl [pH 8.0], 1 mM K_2_HPO_4_/KH_2_PO_4_ [pH 7.6], 8 mM MgSO_4_) or on 1.5% agar supplemented with 1% CTT and kanamycin (50 μg mL^−1^) or oxytetracycline (10 μg mL^−1^) when appropriate (88). Gene expression from the vanillate-inducible promoter (89) was induced with 7.5 µM vanillate. Plasmids were propagated in *E. coli* NEB Turbo at 37°C in lysogeny broth (LB) (90) supplemented with kanamycin (50 μg mL^−1^) or tetracycline (20 μg mL^−1^) when required. *S. enterica* was cultured at 37^°^C in LB supplemented with ampicillin (100 µg mL^-1^), chloramphenicol (30 µg mL^-1^), and 0.2% or 0.4% (w/v) L-arabinose when appropriate.

### Immunoblot analysis

For *M. xanthus*, immunoblot analyses were performed as described (91). Samples were prepared by harvesting *M. xanthus* cells from exponentially growing suspension cultures, followed by resuspension of the cell pellet in SDS lysis buffer (60 mM Tris-HCl [pH 6.8], 2% SDS [w/vol], 10% glycerol [vol/vol], 5 mM ethylenediaminetetraacetic acid, 0.1 M dithiothreitol, 0.005% bromophenol blue [w/vol]). Proteins of an equal amount of cells were loaded per sample. Blots were probed with rabbit polyclonal α-FLAG (1:5,000; Proteintech), mouse monoclonal α-His_6_-tag (1:2000, Proteintech), and rabbit α-PilC (1:2,000) (92) primary antibodies. Rabbit α-FLAG and α-PilC antibodies were used together with horseradish peroxidase (HRP)-conjugated α-rabbit immunoglobulin G (1:15,000; Sigma) as the secondary antibody. Mouse α-His_6_-tag antibodies were used together with HRP-conjugated α-mouse immunoglobulin G (1:2000; GE Healthcare) as the secondary antibody. For detecting biotinylated proteins, blots were probed with Strep-Tactin-HRP conjugate (1:5000; IBA Lifesciences). Blots were developed using Immobilon Forte Western HRP substrate (Millipore) and imaged using the luminescent imager analyzer LAS-4000 (Fujifilm).

For *S. enterica,* cells were grown at 37°C overnight on LB plates supplemented with antibiotics and 0.0%, 0.2% arabinose or 0.4% arabinose. Biomass was collected from each plate, resuspended in 5 mL phosphate-buffered saline (PBS) (137 mM NaCl, 2.7 mM KCl, 10 mM Na_2_HPO_4_, 1.8 mM KH_2_PO_4_, [pH 7.2]), and the optical density at 600 nm adjusted to 5. 1 mL of normalized suspensions was transferred to a microcentrifuge tube and cells harvested by centrifugation (8,000× *g*, 3 min, RT). Cells were resuspended in 1× Laemmli sample buffer (69.4 mM Tris-HCl [pH 6.8], 1.1% (w/vol) SDS, 355 mM β-mercaptoethanol, 11.1% glycerol (vol/vol), 0.005% bromophenol blue (w/vol) to generate whole cell lysates. Samples were incubated 5 min at 65°C, and proteins from an equal number of cells loaded per sample. Proteins were separated by SDS-PAGE on 12% Mini-PROTEAN^®^ TGX™ Precast Protein Gels (Bio-Rad), transferred to a nitrocellulose membrane and blots probed with mouse α-FLAG M2 (Sigma) (1:10,000) and α-DnaK antibodies (Novus Biologicals) (1:5,000) primary antibodies, which were detected using IRDye 680CW-conjugated goat α-mouse IgG (H + L) (LI-COR) and IRDye 800CW-conjugated goat α-mouse IgM, respectively and imaged using a LI-COR Odyssey infrared imaging system.

### LPS O-antigen detection

LPS was extracted from *S. enterica* as described (93, 94). Briefly, cells were grown and harvested as described for *S. enterica* immunoblots. To lyse cells, cell pellets were resuspended in 150 µL of lysis buffer (0.5 M of Tris-HCl [pH 6.8], 2% (w/vol) SDS, 4% β-mercaptoethanol (vol/vol)), and boiled for 10 min. Then, 10 µL of Proteinase K (10 mg mL^−1^) was added and samples incubated at 60°C overnight. To remove proteins, 150 µL of pre-warmed (70°C) 90% phenol solution was added and extracts incubated at 70°C for 15 min with vortexing every 5 min. Then samples were incubated 10 min on ice and centrifuged at 10,000×*g* for 10 min. Finally, 50 µL of the clear aqueous phase was transferred to a clean microcentrifuge tube and 3× loading buffer (0.187 M Tris-HCl [pH 6.8], 6% (w/vol) SDS, 15% (vol/vol) β-mercaptoethanol, 30% (vol/vol) glycerol, 0.03% (w/vol) bromophenol blue) added. LPS samples were separated on 14% (w/vol) Tricine-SDS-PAGE and stained with silver nitrate (93). For immunoblot detection of *Salmonella* O-antigen, LPS samples were separated on 12% Mini-PROTEAN^®^ TGX™ Precast Protein Gels (BioRad), transferred to a nitrocellulose membrane, and probed with rabbit *Salmonella* O antiserum group B (Difco, Beckton Dickinson ref. number 229481) (1:250) together with IRDye 680CW goat α-rabbit immunoglobulin G (1:10,000) (LI-COR) as secondary antibody and detected with a LI-COR Odyssey infrared imaging system (LI-COR).

### Proximity labeling experiments

Protein interactor identification using proximity labeling and shotgun proteomics were done as described (65). Briefly, 25 µM biotin was added to exponentially growing *M. xanthus* strains expressing mTurbo-fused proteins in suspension cultures, and growth allowed to proceed for 3 h. Cells were harvested at 8,000× *g* for 5 min at RT and washed thrice in 5 mL TPM (10 mM Tris-HCl [pH 7.6], 1 mM potassium phosphate [pH 7.6], 8 mM MgSO_4_). The cells were then lysed in 0.6 mL RIPA buffer (50 mM Tris-HCl [pH 7], 150 mM NaCl, 1% Triton X-100 [w/vol], 0.5% sodium deoxycholate [w/vol], 0.2% SDS [w/vol]) also containing 2× protease inhibitor cocktail (Roche) and 5 µL benzonase (Merck). Intact cells were sedimented, and the supernatant was applied to equilibrated G25 desalting columns (GE Healthcare). Next, the protein concentration in the desalted lysate was measured using BCA assay (Thermo Fisher Scientific), and protein concentration was adjusted to 2 mg mL^-1^ in all samples. 50 µL equilibrated magnetic Pierce Streptavidin beads (Thermo Fisher Scientific) were added to the desalted lysate, and the mixture was incubated for 1 h at 4°C with mild shaking. Next, the beads were harvested and washed twice in RIPA, twice in 1 M KCl, and thrice in 50 mM Tris-HCl [pH 7.6]. For the MS analysis of peptides, see Supplementary Materials and Methods.

### Statistical analysis

Data shown for EPS assays, T4P-dependent motility, and immunoblots were obtained in three independent experiments with similar results.

## Author Contributions

Conceived and designed experiments: JS, JM-F, TG, LS-A, MAV

Performed experiments: JS, JM-F, TG

Analyzed data: JS, JM-F, TG, PB, OR, AG, MAV, LS-A

Wrote paper: JS, JM-F, TG, PB, OR, MAV, LS-A

## Notes

### Competing Interest Statement

The authors have declared no competing interest.

