## Supplementary material for "A single-domain response regulator activates exopolysaccharide synthesis by interacting with the initiating phosphoglycosyl transferase": All supplementary information

**This file contains:**

- Supplementary Materials and Methods
- Supplementary Figures 1-12
- Supplementary Tables 1-10
- Supplementary References

**Supplementary Materials and Methods**

Plasmid construction.

For pMP036 (for generating the in-frame deletion in *epsW*), up- and downstream fragments were amplified from genomic DNA of DK1622 using the primer pairs 7420-A/7420-B and 7420-C/7420-D, respectively. Subsequently, the AB and CD fragments were used as templates for an overlapping PCR with the primer pair 7420-A/7420-B to generate the AD fragment. The AD fragment was digested with KpnI/XbaI and cloned into pBJ114.

For pDJS102 (for generating the in-frame deletion in *difE*), up- and downstream fragments were amplified from genomic DNA of DK1622 using the primer pairs 6692-A/6692-B and 6692-C/6692-D, respectively. Subsequently, the AB and CD fragments were used as templates for an overlapping PCR with the primer pair 6692-A/6692-D to generate the AD fragment. The AD fragment was digested with EcoRI/XbaI and cloned in pBJ114.

For pJSc002 (for generating the in-frame deletion in *difD*), up- and downstream fragments were amplified from genomic DNA using the primer pairs 6693-A/6693-B and 6693-C/6693-D, respectively. Subsequently, the AB and CD fragments were used as templates for an overlapping PCR with the primer pair 6693-A/6693-D to generate the AD fragment. The AD fragment was digested with KpnI/XbaI and cloned in pBJ114.

For pJSc003 (for generation of in-frame deletion in *difG*), up- and downstream fragments were amplified from genomic DNA using the primer pairs 6691_A/6691_B and 6691_C/6691_D, respectively. Subsequently, the AB and CD fragments were used as templates for an overlapping PCR with the primer pair 6691_A/6691_D to generate the AD fragment. The AD fragment was digested with KpnI/XbaI and cloned in pBJ114.

For pMP145 (plasmid for expression of $P_{pilA}$ *epsW* from the Mx8 *attB* site): the *epsW* fragment was amplified with the primer pair 7420_PpilA fw /7420_PpilA rev from genomic DNA of DK1622, digested with XbaI/HindIII and cloned into pSW105.

For pJSc113 (for generation of a strain ectopically expressing from the Mx8 *attB* site *epsW* from the *pilA* promoter), a fragment was cleaved from pMP145 using EcoRI/HindIII. The resulting fragment was cloned into pSWU30.

For pJSc105 (for generation of a strain ectopically expressing from the Mx8 *attB* site *mTurbo-epsW-FLAG* from the *pilA* promoter), fragment one was generated from pMH97 (1) using mTurbo-$P_{pilA}$-for and mTurbo-epsW-FLAG int rev. Fragment two was generated from genomic DNA using mTurbo-epsW-FLAG int for and EpsW-FLAG-rev. An overlap extension PCR using mTurbo-$P_{pilA}$-for and EpsW-FLAG-rev was performed on fragments one and two. The resulting fragment was digested with XbaI/HindIII and cloned into pSW105.

For pJSc143 (plasmid for expression of *epsW-His*$_{10}$ from the arabinose-inducible promoter), the *epsW-His*$_{10}$ fragment was amplified with the primer pair 7420_PpilA fw / EpsW-His-rev-HindIII from genomic DNA of DK1622, digested with XbaI/HindIII and cloned into pBAD33.

For pJSc146 (plasmid for expression of $P_{pilA}$ *epsW-His*$_{10}$ from the Mx8 *attB* site), the *epsW-His*$_{10}$ fragment was amplified with the primer pair 7420_PpilA fw / EpsW-His-rev-HindIII from genomic DNA of DK1622, digested with XbaI/HindIII and cloned into pSW105.

Detection of EPS biosynthesis. Colony-based colorimetric EPS assays were performed as described (2). Briefly, cells grown exponentially in suspension cultures were sedimented by centrifugation (3 min, 6,000× *g* at room temperature (RT)) and resuspended in 1% CTT to a calculated cell density of $7 \times 10^9$ cells $mL^{-1}$. Next, 20 µL of the cell suspension was spotted onto 0.5% agar plates containing 0.5% CTT supplemented with either 10 µg $mL^{-1}$ Trypan blue or 20 µg $mL^{-1}$ Congo red. Plates were incubated at 32°C and imaged after 24 h.

Motility assays. Motility assays were conducted as described (3). Briefly, cells grown exponentially in suspension cultures were sedimented by centrifugation (3 min, 6,000× *g*, RT) and resuspended in 1% CTT medium to a calculated cell density of $7 \times 10^9$ cells $mL^{-1}$. Next, 5 µL aliquots of this cell suspension were placed onto 0.5% agar supplemented with 0.5% CTT, followed by incubation at 32°C for 24 h. Imaging was performed using a M205FA stereomicroscope (Leica Microsystems) equipped with a Hamamatsu ORCA-Flash V2 digital CMOS camera (Hamamatsu Photonics).

RNA-seq analysis. Total RNA from exponentially growing *M. xanthus* cells in 20 mL suspension culture was extracted using the Monarch® Total RNA Miniprep Kit (New England Biolabs) according to the manufacturer's protocol. Next, Turbo DNase (Thermo Fisher Scientific) was added to the RNA following the manufacturer's protocol and subsequently removed using the Monarch RNA Cleanup Kit (50 µg; New England Biolabs). Analysis of RNA integrity, ribosomal RNA (rRNA) depletion, library preparation and sequencing was performed by Vertis Biotechnologie. For all samples, RNA integrity was analyzed by capillary electrophoresis using the MultiNA microchip electrophoresis system (Shimadzu). For removal of rRNA, an in-house protocol was used. Briefly, a mixture of *in vitro* transcribed biotin-labelled RNA probes directed against 5S, 16S and 23S bacterial rRNA were mixed with the total RNA samples. The hybridized rRNA molecules were then removed using streptavidin magnetic beads. Next, ribodepleted-RNA samples were fragmented using ultrasound (2 pulses of 30 s each at 4°C). Then, an oligonucleotide adapter was ligated to the 3' end of the RNA molecules. First-strand cDNA synthesis was performed using M-MLV reverse transcriptase and the 3’ adapter as primer. The first-strand cDNA was purified, and the 5' Illumina TruSeq sequencing adapter was ligated to the 3' end of the antisense cDNA. The resulting cDNA was PCR-amplified to about 10−20 ng $\mu L^{-1}$ using a high fidelity DNA polymerase (14 PCR cycles). The cDNA was purified using an Agencourt AMPure XPkit (Beckman Coulter Genomics). For Illumina NextSeq sequencing, the samples were pooled in approximately equimolar amounts. The cDNA pool was purified using the Agencourt AMPure XP kit (Beckman Coulter Genomics). At all steps, quality was assessed *via* the MultiNA microchip electrophoresis system (Shimadzu). The cDNA pool was sequenced on an Illumina NextSeq 500 system using 1× 75 bp read length. All samples had between ~15 and ~20 Mio reads.

The differential gene expression analysis was performed using the RNA-seq pipeline Curare 0.4.4 (4). The reads were preprocessed with Trim Galore (5), trimming low quality ends (<20) and discarding reads smaller than 50 bp. Preprocessed reads were then aligned using Bowtie2 2.4.5 in “very-sensitive” mode and with “–mm” option (6). The resulting alignment files were processed with SAMtools 1.15.1 (7). The subsequent assignment of mapped reads to genome features was done using the featureCounts of the subread 2.0.1 package (8). featureCounts was run with “-s 1” settings assigning reads in a strand-specific manner to the “CDS” features. The differential gene expression was determined with DESeq2 1.34.0 (9). The genome and annotation of *M. xanthus* DK1622 (ASM1268v1) was used for all analyses.

The count table and mapping results have been deposited at EBI ArrayExpress under accession number E-MTAB-14794.

<u>Proteomic analysis using data independent acquisition-mass spectrometry (DIA-MS)</u>. Whole-cell proteomics experiments were performed based on (10). Briefly, *M. xanthus* cells from 10 mL of an exponentially growing suspension culture were harvested by centrifugation (3 min, 10,000 *g*, RT) and washed twice in 0.5 mL PBS (137 mM NaCl, 2.7 mM KCl, 10 mM $Na_2HPO_4$, 1.8 mM $KH_2PO_4$ [pH 7.5] supplemented with 2× protease inhibitor (Roche). The cells were harvested by centrifugation and resuspended in 0.2 mL 0.1 M ammonium bicarbonate containing 2% (w/vol) sodium lauroyl sarcosinate (SLS) and incubated at 95°C for 1 h. Next, the samples were

centrifuged at 14,000× *g* for 5 min, and the supernatant harvested. Next, 1.2 mL freezer-cold acetone was added to the supernatant, mixed, and incubated at −80°C for at least 2 h. Samples were centrifuged at 21,000× *g* for 15 min at 4°C and the supernatant discarded. The pellet was washed thrice with freezer-cold methanol, dried, and the methanol completely removed. The precipitated protein pellets were solubilized using 200 µL 0.5% SLS and heat exposure. The protein amount was determined by bicinchoninic acid (BCA) assay (Thermo Fisher Scientific). Next, proteins were reduced using 5 mM Tris (2-carboxyethyl) phosphine (TCEP, Thermo Fisher Scientific) at 90°C for 15 min and alkylated with 10 mM iodoacetamide (Sigma Aldrich) at 25°C for 30 min in the dark. 50 µg protein was digested using 1 µg trypsin (Serva) at 30°C overnight.

*M. xanthus* peptide samples were acidified to precipitate and remove SLS. Next, the peptides were desalted using C18 solid phase extraction cartridges (Macherey-Nagel), which were prepared for loading the peptide sample by first adding acetonitrile (ACN), followed by 0.1% trifluoroacetic acid (TFA, Thermo Fisher Scientific). The cartridges were equilibrated, peptides were loaded and washed with buffer containing 5% ACN/0.1% TFA. Peptides were eluted with 50% ACN and 0.1% TFA. Dried peptides were reconstituted in 0.1% TFA and then analyzed using liquid-chromatography-mass spectrometry (LC-MS) executed on an Ultimate 3000 RSLCnano connected to an Exploris 480 Mass Spectrometer (all Thermo Fisher Scientific) and a HPLC C18 column packed in-house (75 µm × 42 cm). *M. xanthus* peptides were separated using the following gradient: 94% solvent A (0.15% formic acid) and 6% solvent B (99.85% acetonitrile, 0.15% formic acid) to 25% solvent B over 95 minutes at a flow rate of 300 nL $min^{-1}$, followed by an additional increase of solvent B 35% over 25 min.

MS raw data was acquired in data-independent acquisition (DIA) mode. Briefly, the spray voltage was set to 2.3 kV, the funnel radio frequency level was set to 40, and the ion transfer capillary was heated to 275 °C. For DIA experiments, full MS resolutions were set to 120,000 at m/z 200 and full MS, with the AGC (automatic gain control) target set to 300% and ion accumulation time (IT) of 50 ms. The AGC target value for fragment spectra was set to 3000%. 45 windows, each set to 14 Da plus 1 Da overlap, were used. The resolution was set to 15,000 with MS/MS IT to 22 ms. Stepped HCD high energy collision dissociation (HCD) collision energies of 25, 27.5, 30% were applied. MS1 data was acquired in profile mode, and MS2 DIA data in centroid mode.

DIA data analysis was performed using the neural network (NN)-based DIA-NN suite version 1.8 (11) in combination with a *M. xanthus* protein database from UniProt. A dataset centric spectral library was generated for the DIA analysis. DIA-NN applied noise interference correction (mass correction, RT prediction and precursor/fragment co-elution correlation) and peptide precursor signal extraction from the raw DIA-NN data. The following settings were used for the analysis: Full tryptic digestion with two missed cleavage sites was allowed, with oxidized methionines and carbamidomethylated cysteines as modifications. “Match between runs” and “remove likely interferences” were enabled. The NN classifier was set to the single-pass mode, and protein inference was based on genes. Quantification was performed using the “any LC (high accuracy)” strategy, with cross-run normalization set to RT-dependent. Library generation was set to smart profiling. Outputs from DIA-NN were further evaluated using a SafeQuant (12, 13) script modified to process DIA-NN results. For global analyses, imputation of missing values was performed using a normal distribution function, similar to the strategy implemented in the Perseus statistical software package (14).

The mass spectrometry proteomics data of whole cell proteomics experiments have been deposited to the ProteomeXchange Consortium (15) *via* the PRIDE (16) partner repository with the dataset identifier PXD063981.

In the proximity labeling experiments, proteins were digested using an on-bead digest with 1 µg trypsin overnight at 30°C in the presence of 1 mM TCEP. Following digestion, the peptides were further alkylated with 2 mM iodoacetamide and peptides purified using solid-phase extraction on Chromabond spin columns (Macherey-Nagel). Peptides were then loaded on the LC-MS system described above. Peptide separation was carried out using a constant flow rate of 300 nL $min^{-1}$ and a 45 min gradient from 6%–35% LC buffer B (99.85% acetonitrile/0.15% formic acid). Eluting peptides were ionized at 2.3 kV. MS raw data was acquired in data-dependent mode (DDA) with a MS1 resolution of 60,000 full width at half maximum (at m/z 200) followed by MS/MS scans of the most intense ions within 1 s (cycle 1 s). The dynamic exclusion time was set to 14 s, and the ion accumulation time was set to 50 ms for MS1 and 50 ms at 17,500 resolution for MS/MS scans. The AGC was set to $3\times10^6$ for MS survey scans and $2\times10^5$ for MS/MS scans. The quadrupole isolation was 1.5 m/z, collision was induced with an HCD collision energy of 27 %.

MS raw data was analyzed with MaxQuant (17), and an *M. xanthus* UniProt database. MaxQuant was used with standard settings without the “match between runs” option. The MaxQuant “proteinGroups.txt” output file was further processed by the SafeQuant R package for statistical analysis (12, 13). The same data imputation method was used as for the global proteome analysis. The criteria for significant enrichment in samples over that of the control were: An abundance difference of ≥8 ($log_2$-fold enrichment of ≥3.0) and a *P*-value of ≤0.0001 ($-log_{10}$ *P*-

value ≥ 4.0). *P*-values were calculated using an eBayes method provided by the Limma software package (18).

Proximity labeling experiment data have been deposited to the ProteomeXchange Consortium (15) *via* the PRIDE (16) partner repository with the dataset identifier PXD063973.

Bioinformatics. Gene and protein sequences were fetched from the KEGG (19) and UniProt (20) databases. The operon structure of the *eps* loci was generated using the RNA-seq and cappable-seq data from (21). The phylogenetic tree of Myxobacteria was generated in MEGA-X (22) using the neighbor-joining method (23) and the genome sequences listed in Table S10. Myxobacterial EpsZ and EpsW orthologs were previously identified (2, 10).

Protein structure predictions were conducted using AlphaFold2 and AlphaFold2-Multimer_v3 implemented *via* ColabFold (24-26) with the Alphafold2_mmseqs2 notebook and default settings. To assess the quality of AlphaFold-generated models, predicted local distance difference test (pLDDT) and predicted alignment error (pAE) graphs of five models were generated using a custom-made Matlab R2020a (The MathWorks) script. These models were assessed based on a combination of pLDDT, pAE values, and the interface predicted template modeling score (ipTM) scores. The best-ranked models were then selected for analysis and presentation. Per-residue model confidence was inferred from pLDDT values (>90, high accuracy; 70 to 90, generally good accuracy; 50 to 70, low accuracy; <50, should not be interpreted) (24). To analyze the relative positioning of residues, the pAE, measured in Å, was assessed. The lower the pAE value, the higher the accuracy of the relative position of residue pairs and, consequently, the relative position of domains/subunits/proteins (24). ipTM values (25) were further used to assess the reliability of interfaces in multimeric models. An ipTM score above 0.80 indicates an accurate interface, scores between 0.60 and 0.80 fall into a grey zone where predictions may or may not be correct, and scores below 0.60 indicate a failed prediction (27, 28). Protein interfaces in multimeric structural models were determined using PDBePISA with default settings (29).

To inspect and visualize structural models, we used PyMOL (The PyMOL Molecular Graphics System, Version 2.4.1 Schrödinger, LLC). Models colored based on pLDDT values were made using a custom command line (spectrum b, red_yellow_green_cyan_blue, minimum=50, maximum=90). Foldseek (30) was used to identify protein homologs in the $PDB^{100}$ database. The positioning of protein structural models within the IM was assessed using the PPM 3.0 web server (31) with default settings and membrane type set to “Gram-negative bacteria inner membrane”. The coordinates of all structural models generated in this study have been deposited in the Edmond research data repository under ref (32).

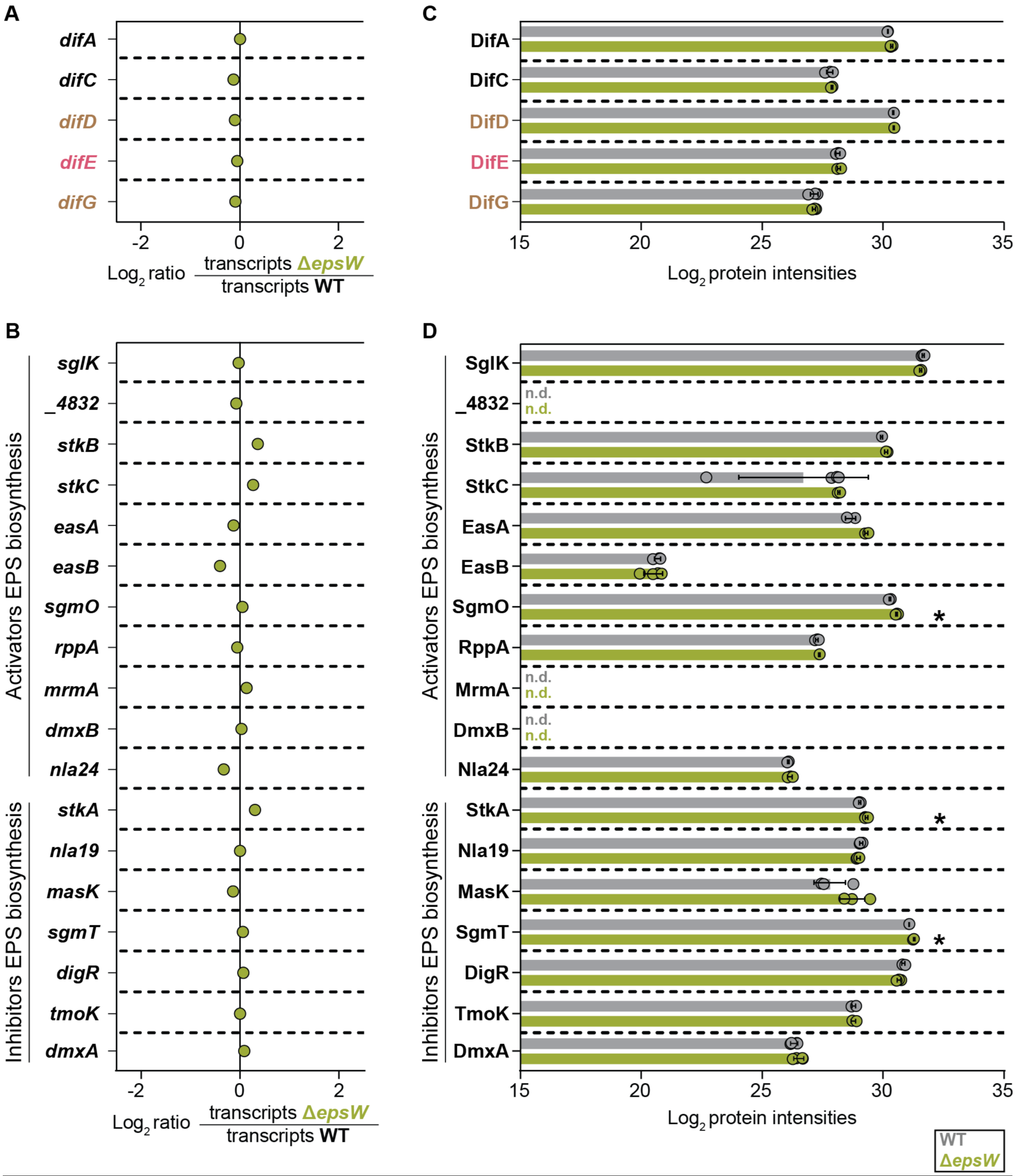


Figure S1. EpsW is neither required for the accumulation of Dif proteins nor of other regulators of EPS biosynthesis.

(A, B) Differential expression of *dif* genes (A) and other genes encoding regulators of EPS biosynthesis (B) in the Δ*epsW* mutant compared to WT. The RNAseq experiment was performed with four biological replicates per strain from cells grown in suspension culture. X-axis, $log_2$-fold ratio of the mean transcripts in the Δ*epsW* mutant over the mean transcripts in the WT calculated using the DESeq2 method (9). Statistical analysis was performed in the DESeq2 analysis. No significant differences were identified (adjusted $P$ value ≤ 0.001). For detailed descriptions of the regulators in B see (33).

(C, D) Protein abundance in whole-cell proteomes of the Δ*epsW* mutant compared to WT. The LFQ-MS-based proteomics was performed with four biological replicates from cells grown as in (A, B). X-axis, normalized $\log_2$ intensities of proteins in the indicated strains. Data points represent each of the four biological replicates. Error bars, mean ± standard deviation (SD) based on these replicates. *, $P < 0.001$ (Welch's test); n. d., not detected.

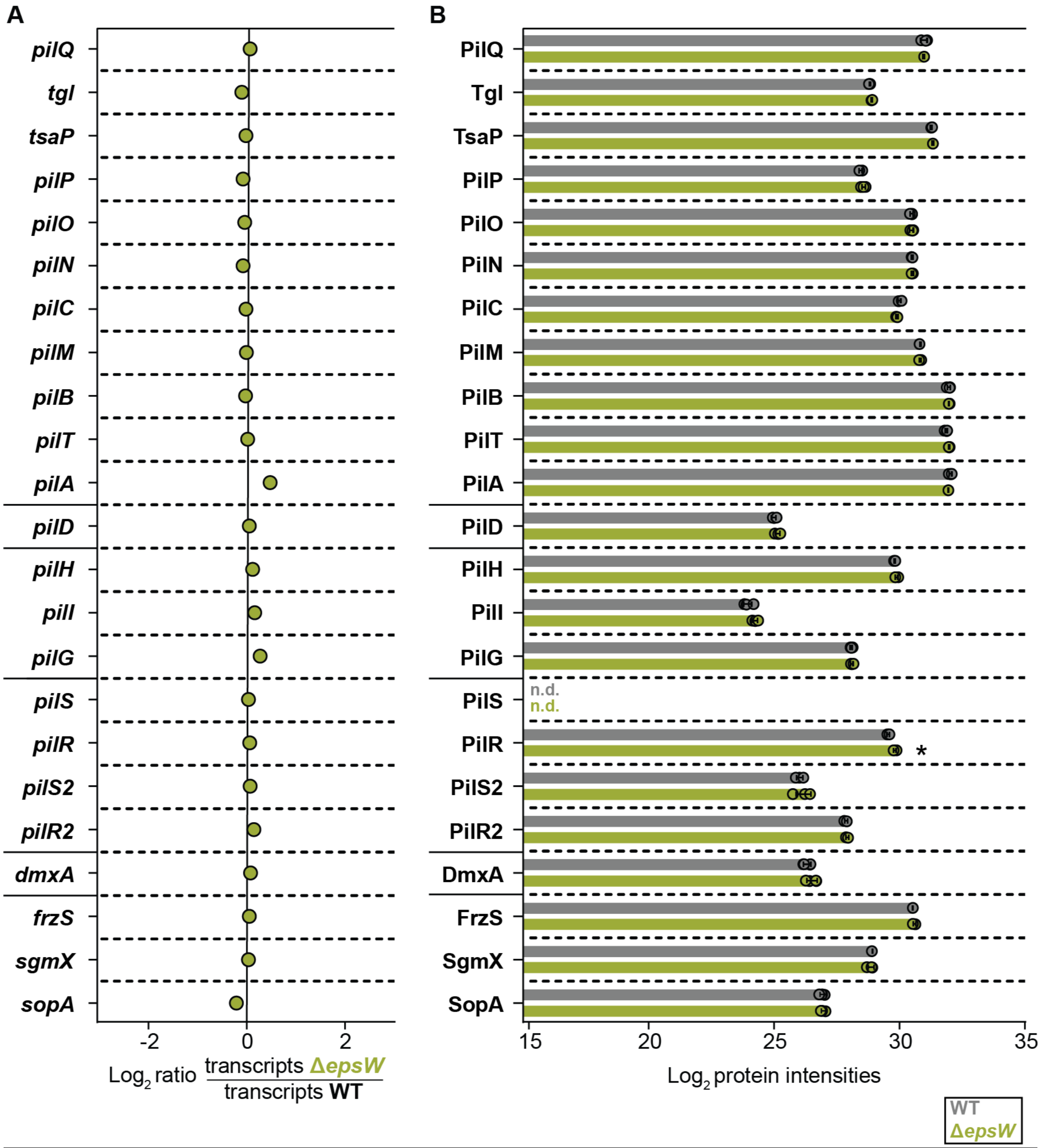


Figure S2. EpsW is not required for the accumulation of proteins required for T4P formation and function.

(A) Differential expression of genes encoding proteins required for T4P formation and function in the Δ*epsW* mutant compared to WT. The RNAseq experiment was performed with four biological replicates per strain from cells grown in suspension culture. X-axis, $log_2$-fold ratio of the mean transcripts in the Δ*epsW* mutant over the mean transcripts in the WT calculated using the DESeq2 method (9). Statistical analysis was performed in the DESeq2 analysis. No significant differences were identified (adjusted *P* value ≤ 0.001). PilQ is the multimeric OM secretin stabilized by LysM-domain protein TsaP (34-37). Tgl stimulates PilQ multimerization (36, 37). PilN/-O/-P are structural components in the periplasm. PilC/-M form the IM/cytoplasmic platform complex. PilB/-T are the extension and retraction ATPases, respectively, and PilA is the major pilin (35, 38). PilH/-I/-G are suggested to form an ABC transporter (39). PilD is the prepilin leader peptidase (39, 40). PilR/-S/-R2/-S2 are regulatory proteins (41, 42). DmxA is the diguanylate cyclase important for stimulating c-di-GMP synthesis during cytokinesis (43). FrzS, SgmX, and SopA jointly regulate T4P formation (44-47).

(B) Protein abundance in whole-cell proteomes of the Δ*epsW* mutant compared to WT. The LFQ-MS-based proteomics was performed with four biological replicates from cells grown as in (A). X-axis, normalized $\log_2$ intensities of proteins in the indicated strains. Data points represent each of the four biological replicates. Error bars, mean ± SD. *, $P < 0.001$ (Welch's test); n.d., not detected.

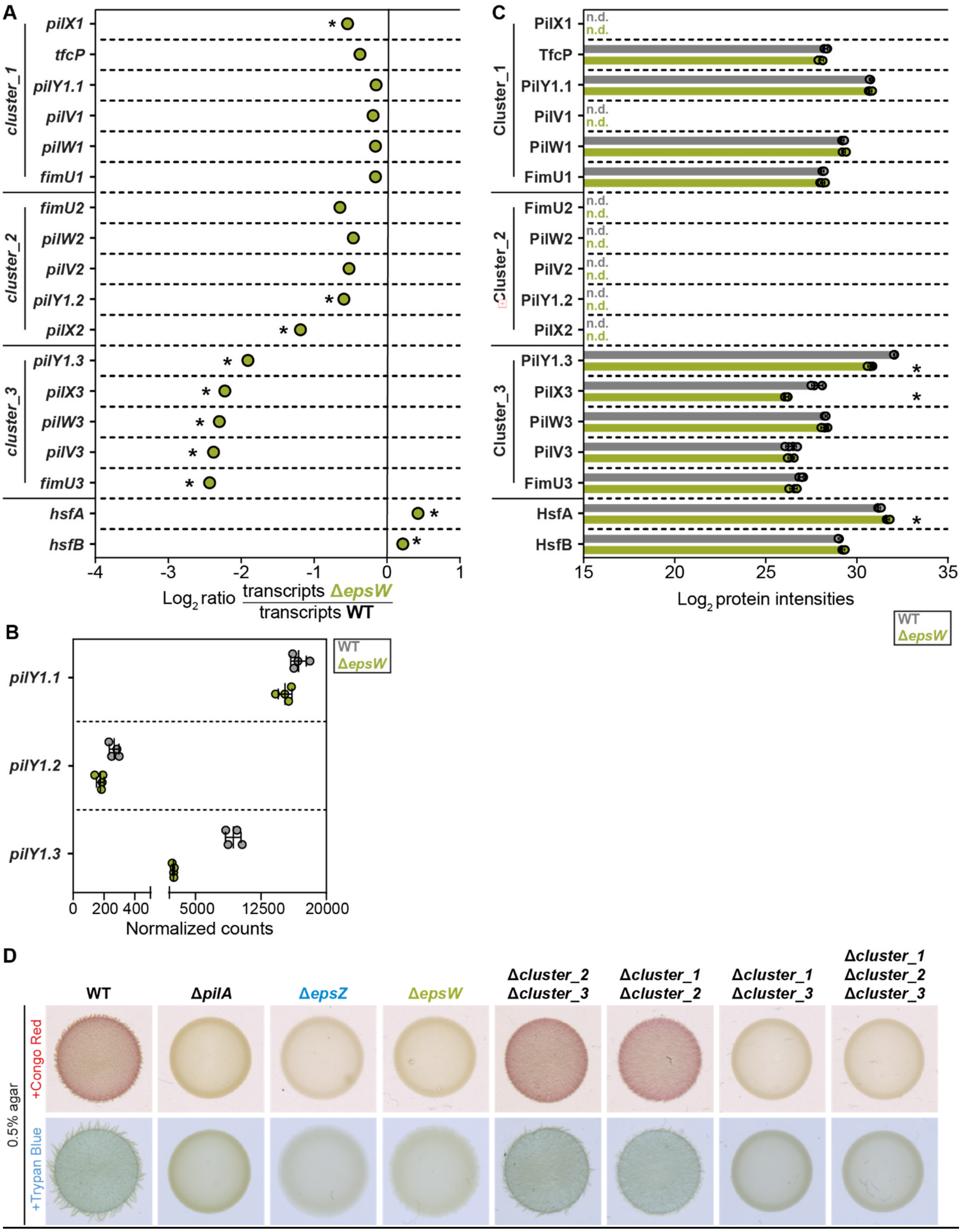


Figure S3. EpsW is important for the accumulation of PilY1.3 and PilX3.

(A) Differential expression of genes encoding cluster_1, cluster_2 and cluster_3 T4P priming complexes in the Δ*epsW* mutant compared to the WT. The RNAseq experiment was performed with four biological replicates per strain from cells grown in suspension culture. X-axis, $\log_2$-fold ratio of the mean transcripts in the Δ*epsW* mutant over the mean transcripts in the WT calculated using the DESeq2 method (9). Statistical analysis was performed in the DESeq2 analysis. *, adjusted *P*-value ≤0.001.
(B) Normalized read counts for *pilY1.1*, *pilY1.2* and *pilY1.3* in the RNAseq experiment of the Δ*epsW* mutant and the WT.
(C) Protein abundance in whole-cell proteomes of the Δ*epsW* mutant compared to WT. The LFQ-MS-based proteomics was performed with four biological replicates from cells grown as in (A). X-axis, normalized $\log_2$ intensities of the indicated proteins in the indicated strains. Data points represent each of the four biological replicates. Error bars, mean ± SD. *, $P < 0.001$ (Welch's test); n.d., not detected.
(D) Cluster_1 encoding minor pilins and PilY1.1 is sufficient for EPS biosynthesis. EPS biosynthesis was assessed by spotting cells on 0.5% agar supplemented with 0.5% CTT and either Congo red or Trypan blue, and images were recorded after 24 h. The Δ*epsZ* mutant was used as the negative control for EPS biosynthesis. For comparison, the Δ*pilA* mutant, which lacks the major pilin of T4P (42), was included. Similarly, different cluster mutants were included, demonstrating that cluster_3 is also sufficient for EPS biosynthesis, while cluster_2 is not.

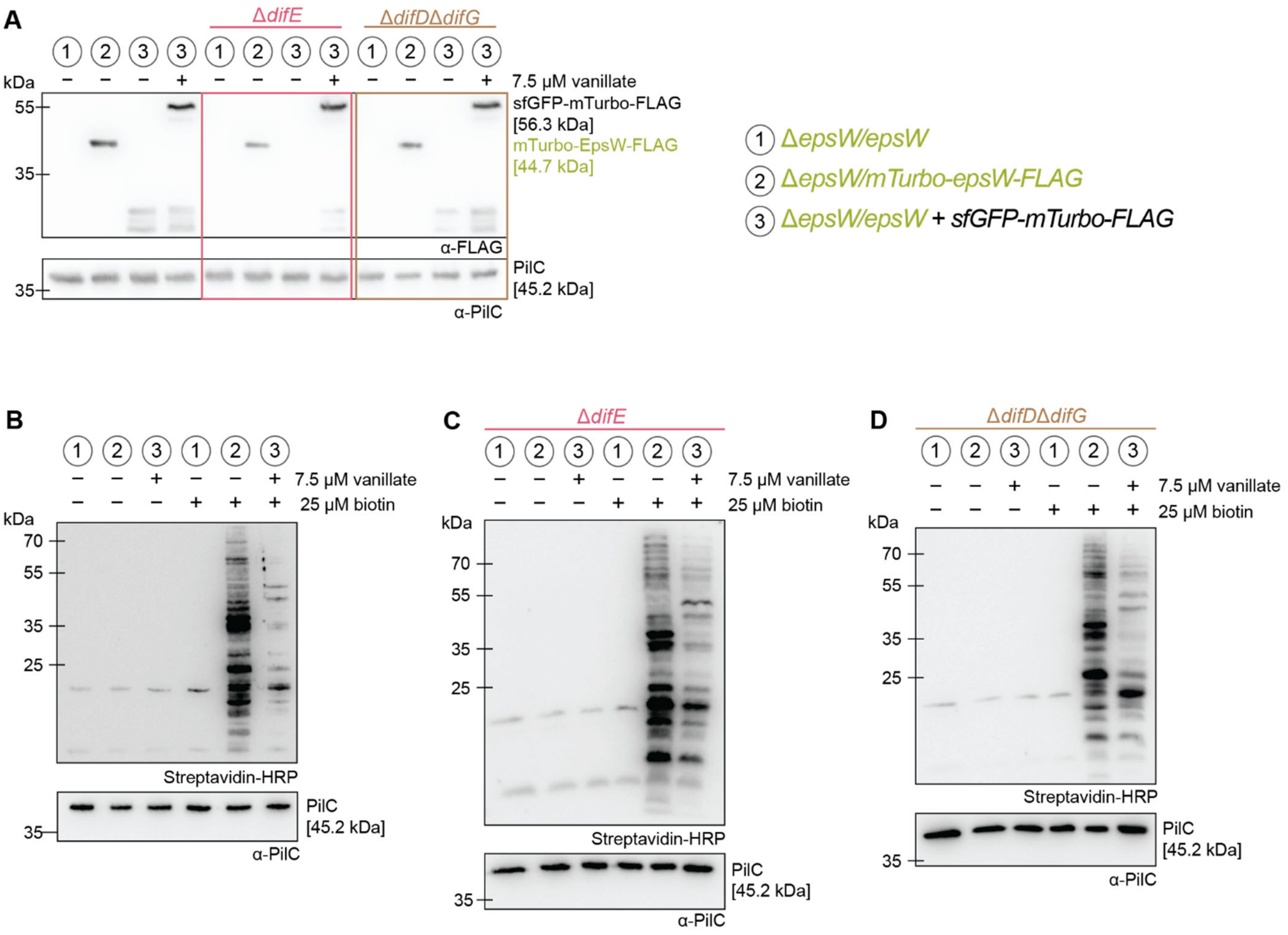


Figure S4. mTurbo-EpsW-FLAG and sfGFP-mTurbo-Flag accumulate and have biotin ligase activity *in vivo*

(A) Immunoblots of mTurbo-EpsW-FLAG and sfGFP-mTurbo-FLAG abundance. Total cell lysates from an equal amount of cells of the indicated strains were separated by SDS-PAGE, followed by immunoblotting with α-FLAG antibodies. The upper blot was stripped and reprobed with α-PilC antibodies as a loading control. +/− symbols indicate whether 7.5 µM vanillate was added or not for 18 h. The code for the different strains is indicated on the right. *epsW* and *mTurbo-epsW-FLAG* were ectopically expressed from the *pilA* promoter from a plasmid integrated in a single copy at the Mx8 *attB* site. *sfGFP-mTurbo-FLAG* was expressed from the vanillate-inducible promoter from a plasmid integrated in a single copy at the *18-19* locus.

(B-D) The mTurbo constructs have biotin ligase activity. Total cell lysates from an equal amount of cells of the indicated strains were analyzed by SDS-PAGE, followed by blotting and detection using Streptavidin-HRP. The blots were subsequently stripped and reprobed with α-PilC antibodies as a loading control. +/− symbols indicate whether 7.5 µM vanillate was added or not for 18 h and whether they were incubated with 25 µM biotin for 3 h. The code for the different strains is as in A. Note that the mTurbo constructs had low biotin-ligase activity when grown without biotin, indicating low endogenous biotinylation of proteins.

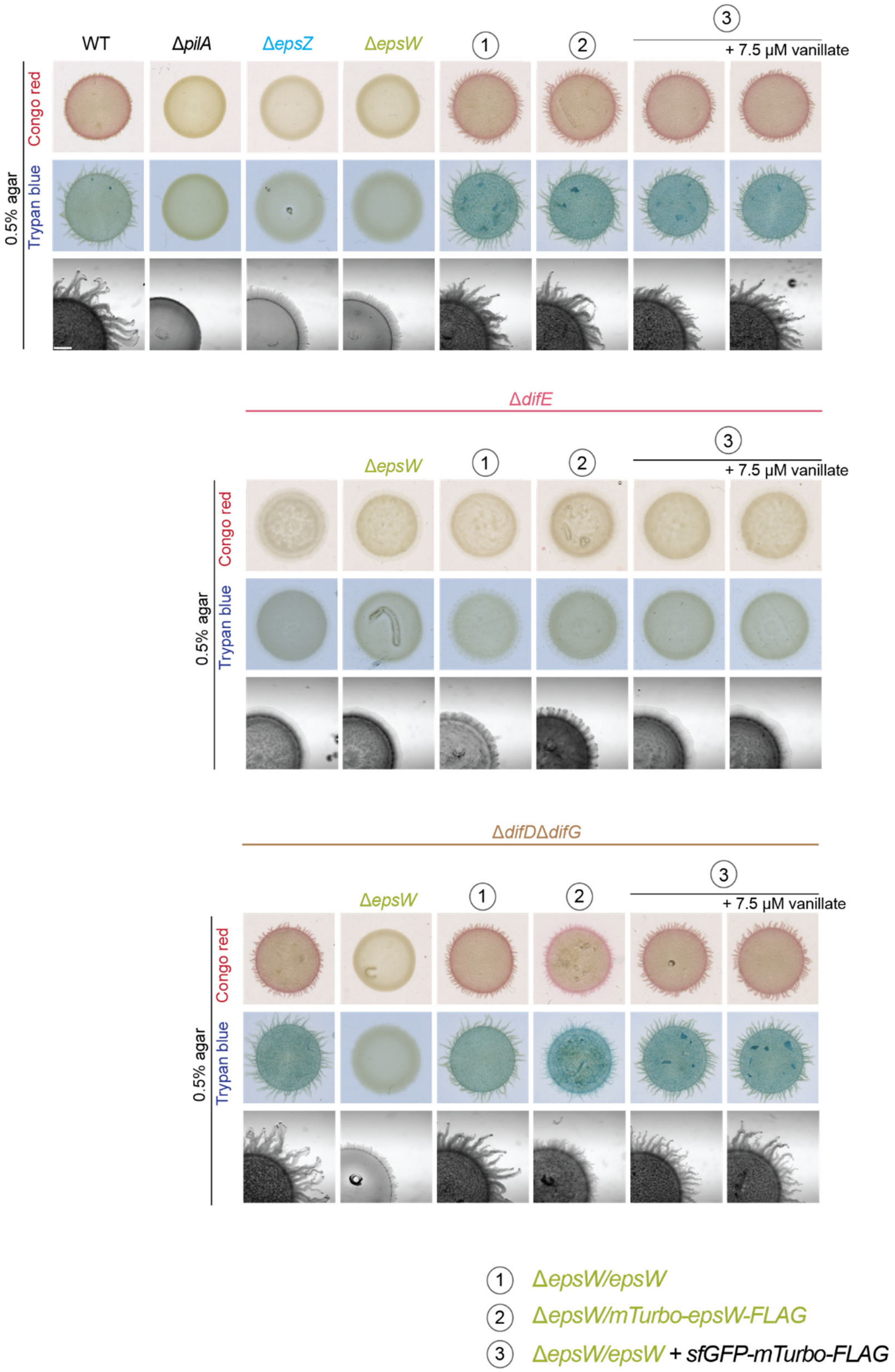


Figure S5. mTurbo-EpsW-FLAG is functional
EPS biosynthesis and T4P-dependent motility in the indicated strains. EPS biosynthesis was assessed on 0.5% agar containing 0.5% CTT and either Congo red or Trypan blue, and images

were captured after 24 h. As the negative control for EPS biosynthesis, the Δ*epsZ* mutant was used. T4P-dependent motility was analyzed on 0.5% agar with 0.5% CTT, with images recorded after 24 h. The Δ*pilA* mutant, which lacks the major subunit of T4P (42), was used as the negative control for T4P-dependent motility. The code for the different strains is indicated below. Scale bar, 1 mm.

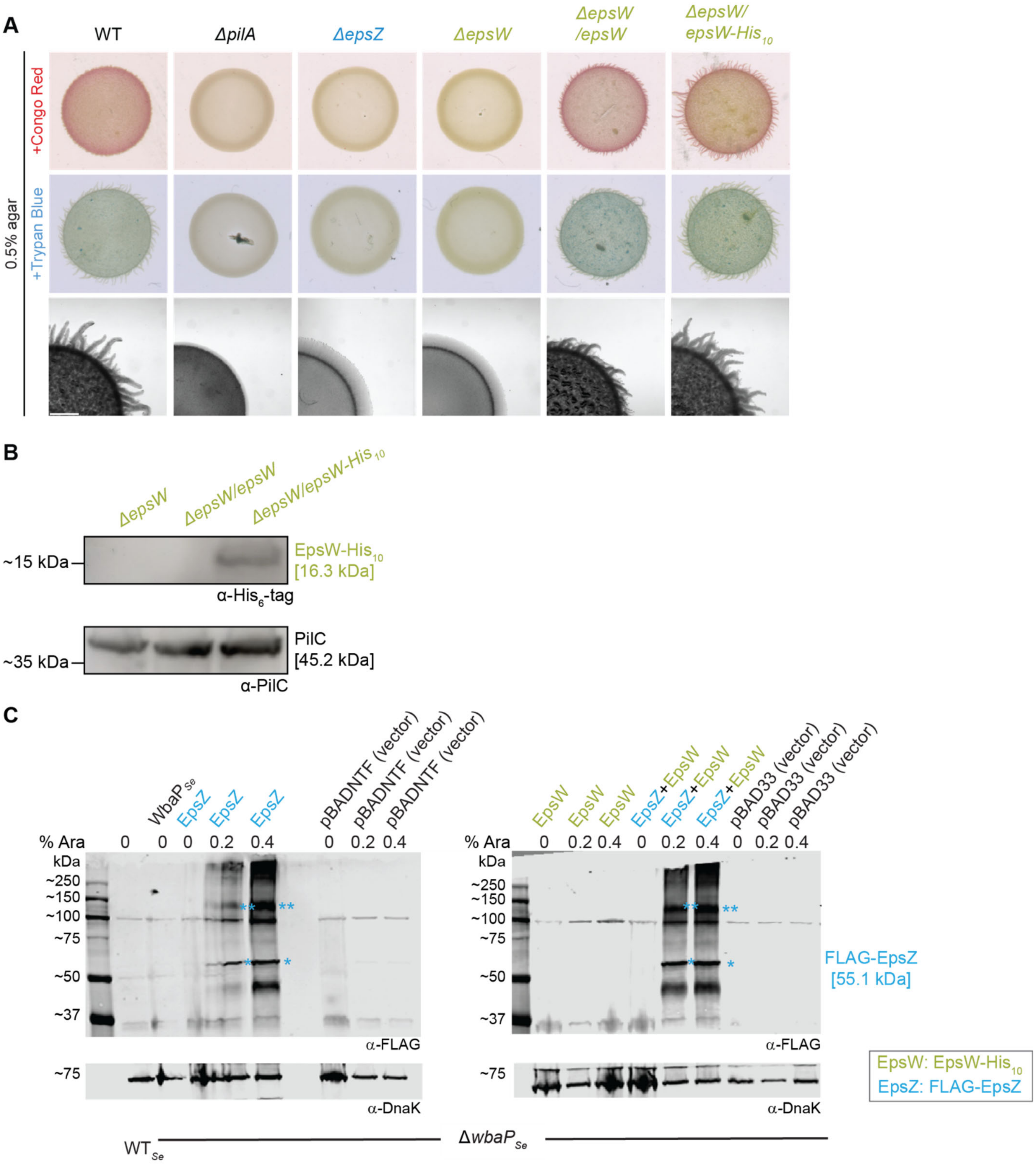


Figure S6. EpsW-His$_{10}$ is functional and accumulates in *M. xanthus* and FLAG-EpsZ accumulates in *S. enterica*.

(A) EPS biosynthesis and T4P-dependent motility in the indicated *M. xanthus* strains. EPS biosynthesis was assessed on 0.5% agar containing 0.5% CTT and either Congo red or Trypan

blue, and images were captured after 24 h. As the negative control for EPS biosynthesis, the Δ*epsZ* mutant was used. T4P-dependent motility was analyzed on 0.5% agar with 0.5% CTT, with images recorded after 24 h. The Δ*pilA* mutant, which lacks the major subunit of T4P (42), was used as the negative control for T4P-dependent motility. In the Δ*epsW/epsW* strain and the Δ*epsW/epsW-His*$_{10}$ strains, the respective genes were ectopically expressed from the *pilA* promoter from plasmids integrated in a single copy at the Mx8 *attB* site. Scale bar, 1 mm.
(B) Immunoblot analysis of EpsW-His$_{10}$. Total cellular lysates from an equal amount of cells of the indicated *M. xanthus* strains were separated *via* SDS-PAGE, followed by immunoblotting with α-His$_6$-tag antibodies. The upper blot was stripped and reprobed with α-PilC antibodies as loading control.
(C) Immunoblot analysis of FLAG-EpsZ abundance in the *S. enterica* Δ*wbaP* mutant (Δ*wbaP*$_{Se}$). The indicated proteins were expressed as in the experiment to detect LPS O-antigen in the presence of arabinose as indicated. Total cellular lysates from an equal amount of cells of the indicated *S. enterica* strains were separated by SDS-PAGE and probed with α-FLAG antibodies and α-DnaK antibodies as a loading control. * and ** indicate the monomeric and oligomeric forms of FLAG-EpsZ. Upper and lower panels are from the blots. All lysates were prepared in the same experiment and separated on by SDS-PAGE on two gels. Note that WbaP of *S. enterica* (WbaP$_{Se}$) is not FLAG-tagged and, therefore, not detected. FLAG-EpsZ and EpsW-His$_{10}$ were expressed from pMP146 (vector: pBADNTF) and pJSc143 (vector pBAD33), respectively under the control of an arabinose-inducible promoter.

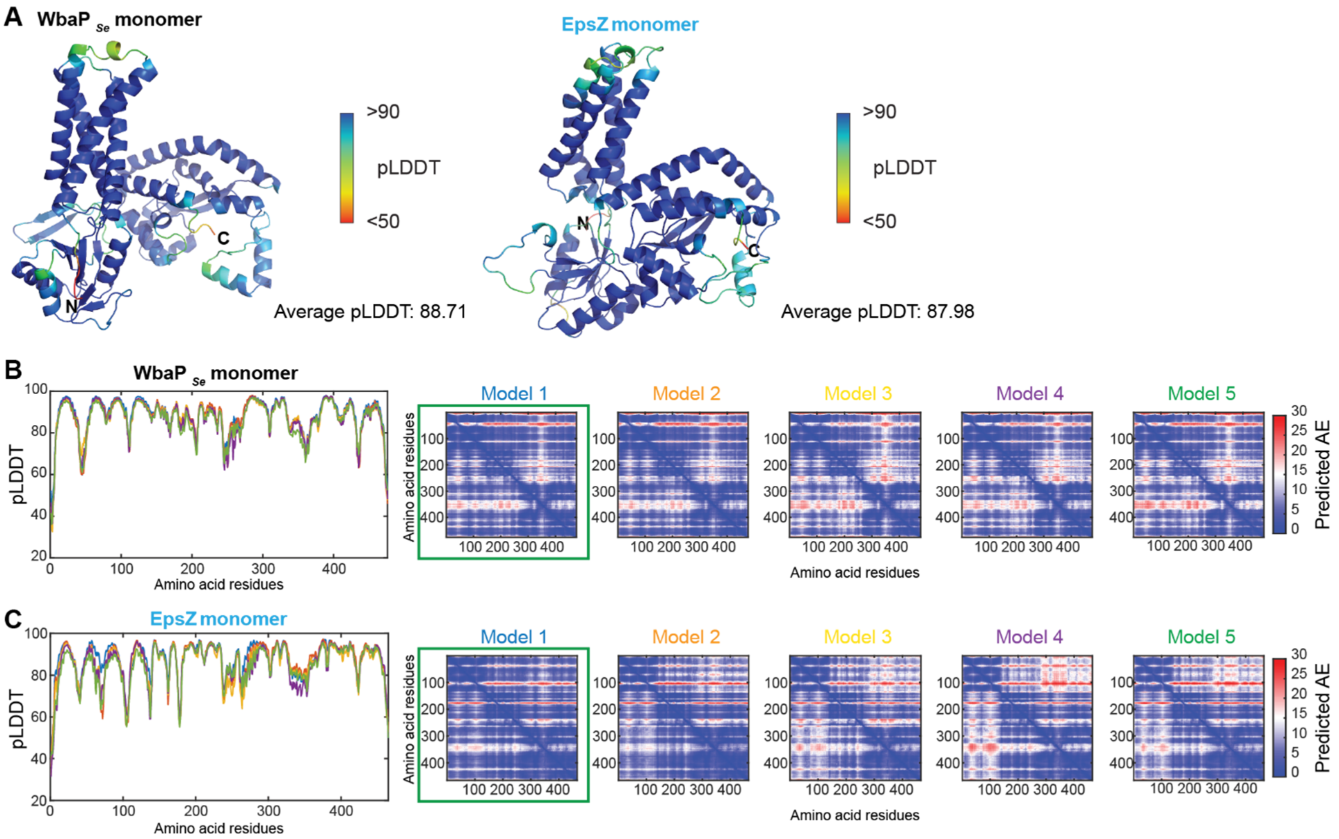


Figure S7. AlphaFold2 models of monomeric $WbaP_{Se}$ and EpsZ.

(A) Left panel, model rank 1 of $WbaP_{Se}$ colored according to pLDDT score, with the average pLDDT indicated. Right panel, model rank 1 of EpsZ colored according to pLDDT score, with the average pLDDT indicated. N- and C-termini are shown.

(B) Left panel, pLDDT plot shown for the five generated models of $WbaP_{Se}$. Right panel, pAE plots shown for the five generated models of $WbaP_{Se}$. Model rank 1 (highlighted by a green box) was selected for further analyses.

(C) Left panel, pLDDT plot shown for the five generated models of EpsZ. Right panel, pAE plots shown for the five generated models of EpsZ. Model rank 1 (highlighted by a green box) was selected for further analyses.

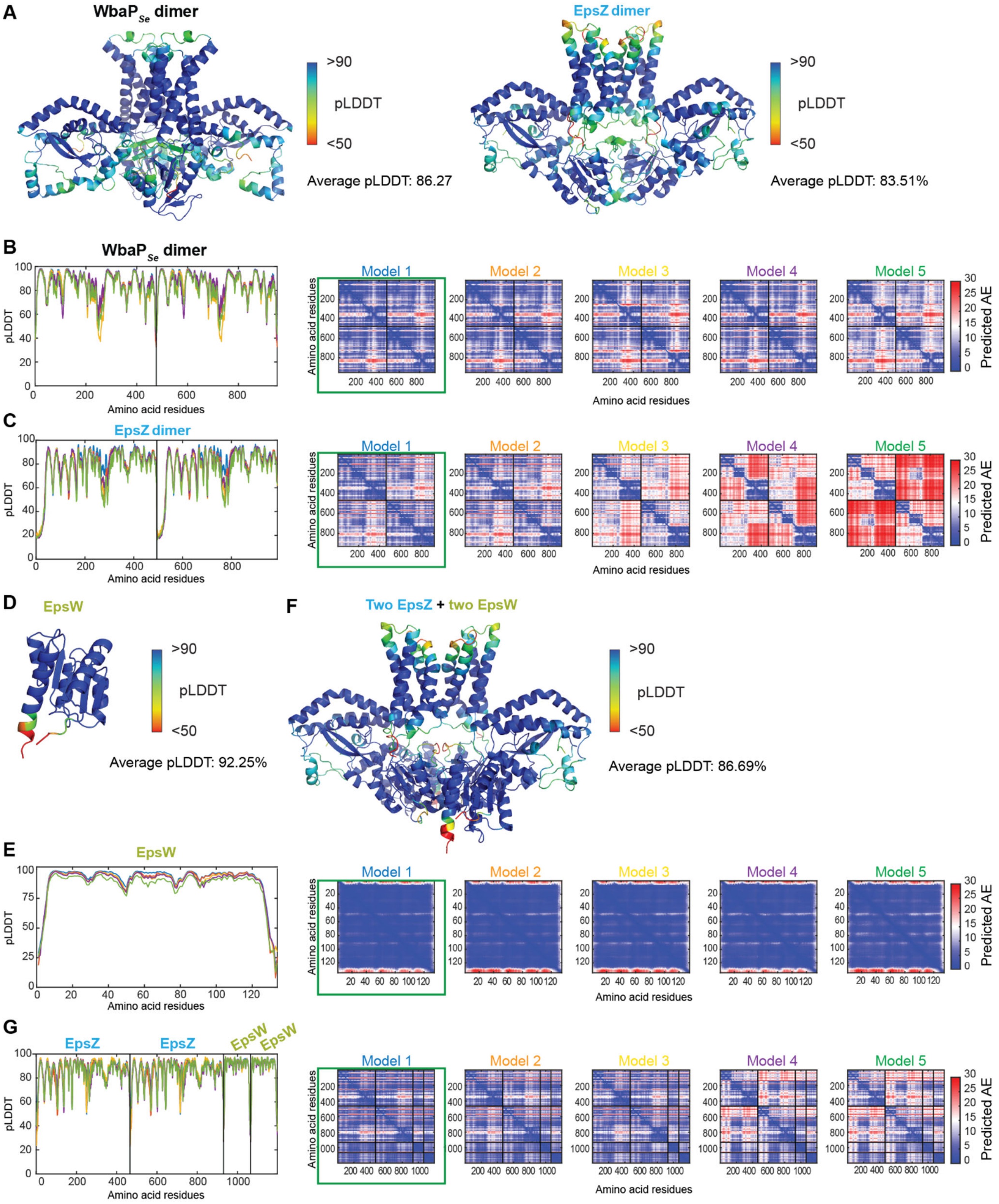


Figure S8. AlphaFold2-Multimer models of the WbaP$_{Se}$ dimer, the EpsZ dimer, and the EpsZ-EpsW heterocomplex in 2:2 stoichiometry.

(A) Left panel, model rank 1 of the $WbaP_{Se}$ dimer colored according to pLDDT score, with average pLDDT indicated. Right panel, model rank 1 of the EpsZ dimer colored according to pLDDT score, with average pLDDT indicated.
(B) Left panel, pLDDT plot shown for the five generated models of the $WbaP_{Se}$ dimer. Right panel, pAE plots shown for the five generated models of the $WbaP_{Se}$ dimer. Model rank 1 (highlighted by a green box) was selected for further analyses.
(C) Left panel, pLDDT plot shown for the five generated models of the EpsZ dimer. Right panel, pAE plots shown for the five generated models of the EpsZ dimer. Model rank 1 (highlighted by a green box) was selected for further analyses.
(D) Model rank 1 of EpsW colored according to pLDDT score, with average pLDDT indicated.
(E) Left panel, pLDDT plot shown for the five generated models of EpsW. Right panel, pAE plots shown for the five generated models of EpsW. Model rank 1 (highlighted by a green box) was selected for further analyses.
(F) Model rank 1 of the EpsZ-EpsW heterocomplex in 2:2 stoichiometry colored according to pLDDT score, with average pLDDT indicated.
(G) Left panel, pLDDT plot shown for the five generated models of the EpsZ-EpsW heterocomplex in 2:2 stoichiometry. Right panel, pAE plots shown for the five generated models of the EpsZ-EpsW heterocomplex in 2:2 stoichiometry. Model rank 1 (highlighted by a green box) was selected for further analyses.

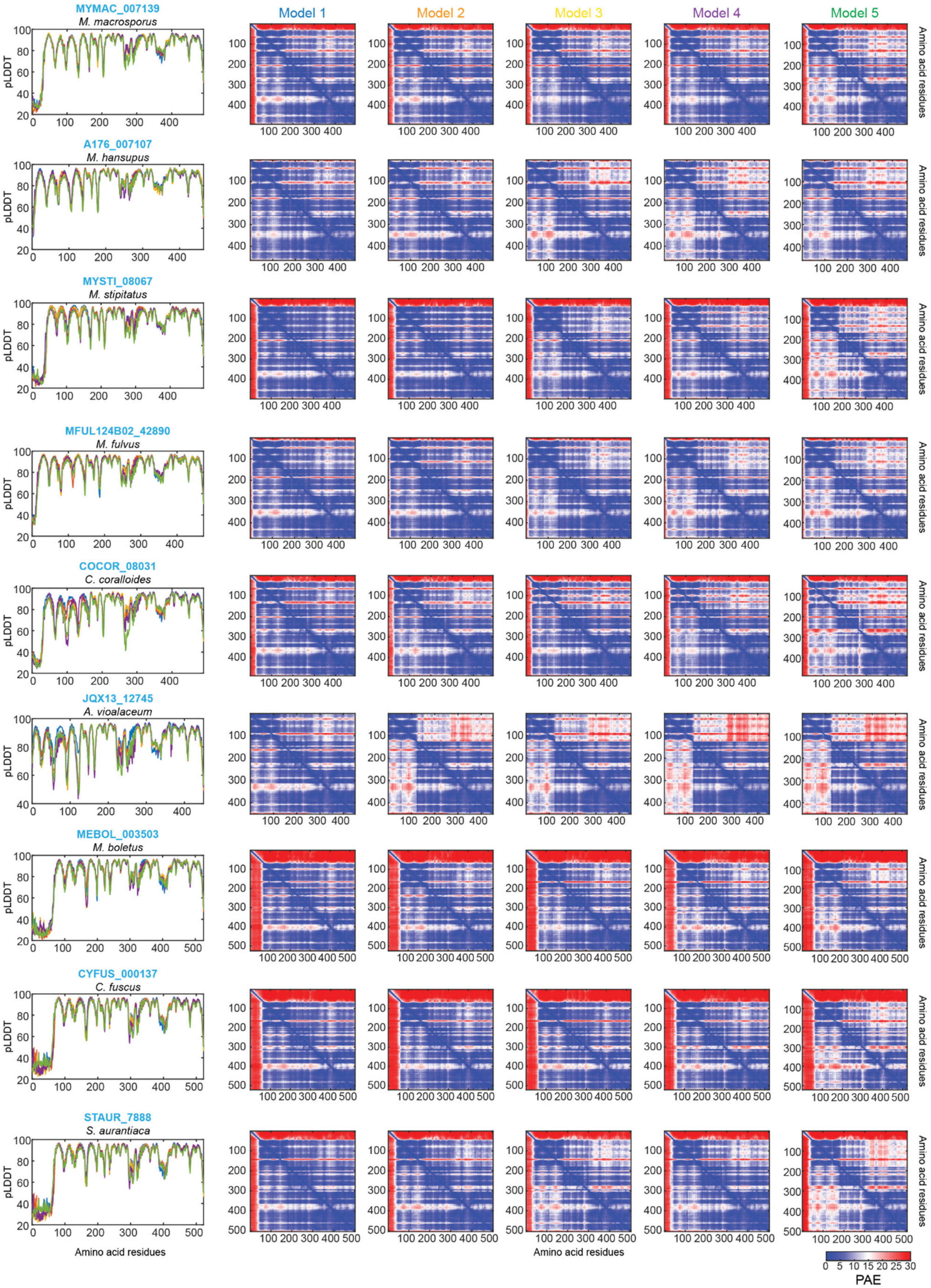
MYMAC_007139
M. macrosporus
A176_007107
M. hansupus
MYSTI_08067
M. stipitatus
MFUL124B02_42890
M. fulvus
COCOR_08031
C. coralloides
JQX13_12745
A. vioalaceum
MEBOL_003503
M. boletus
CYFUS_000137
C. fuscus
STAUR_7888
S. aurantiaca
pLDDT
Amino acid residues
Model 1
Model 2
Model 3
Model 4
Model 5
Amino acid residues
PAE
0 5 10 15 20 25 30

Figure S9. AlphaFold2 models of monomers of nine myxobacterial EpsZ orthologs encoded together with an EpsW ortholog.

Left panels, pLDDT plot shown for the five generated models of each EpsZ ortholog. Right panels, pAE plots shown for the five generated models of each EpsZ ortholog. Model rank 1 were selected for further analyses. Of note, in some models, the N-terminus is unstructured and has low confidence.

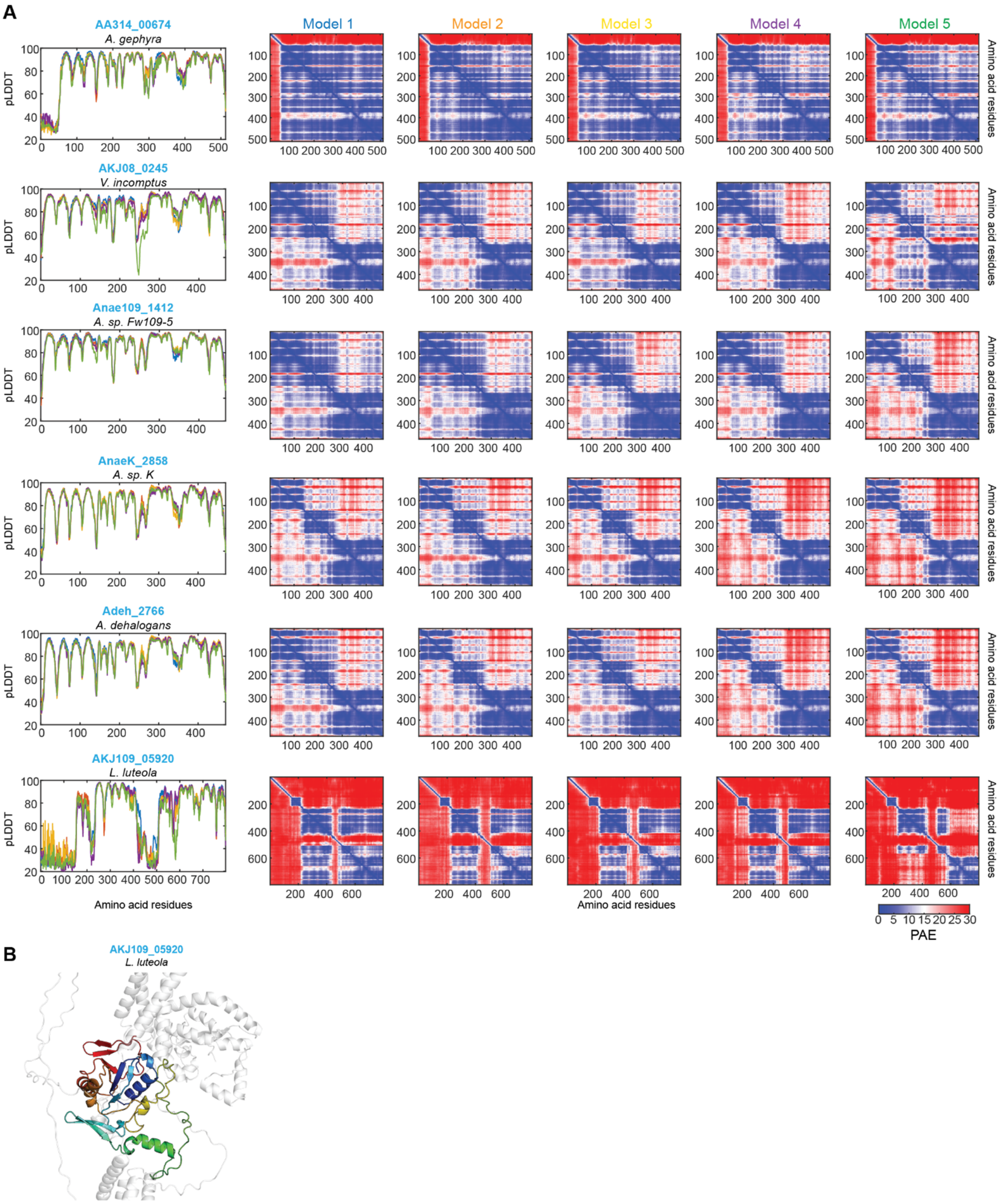


Figure 10. AlphaFold2 models of six myxobacterial EpsZ orthologs encoded without an EpsW ortholog.

(A) Left panels, pLDDT plot shown for the five generated models of each EpsZ ortholog. Right panels, pAE plots shown for the five generated models of each EpsZ ortholog. Model rank 1 were

selected for further analyses. Of note, in some models, the N-terminus is unstructured and has low confidence.
(B) Computational structural model of the EpsZ ortholog AKJ109_05920 in *Labilithrix luteola*. The cytoplasmic DUF of the rank 1 AlphaFold2 model is shown and depicted using a gradient from blue (N-terminus) to red (C-terminus).

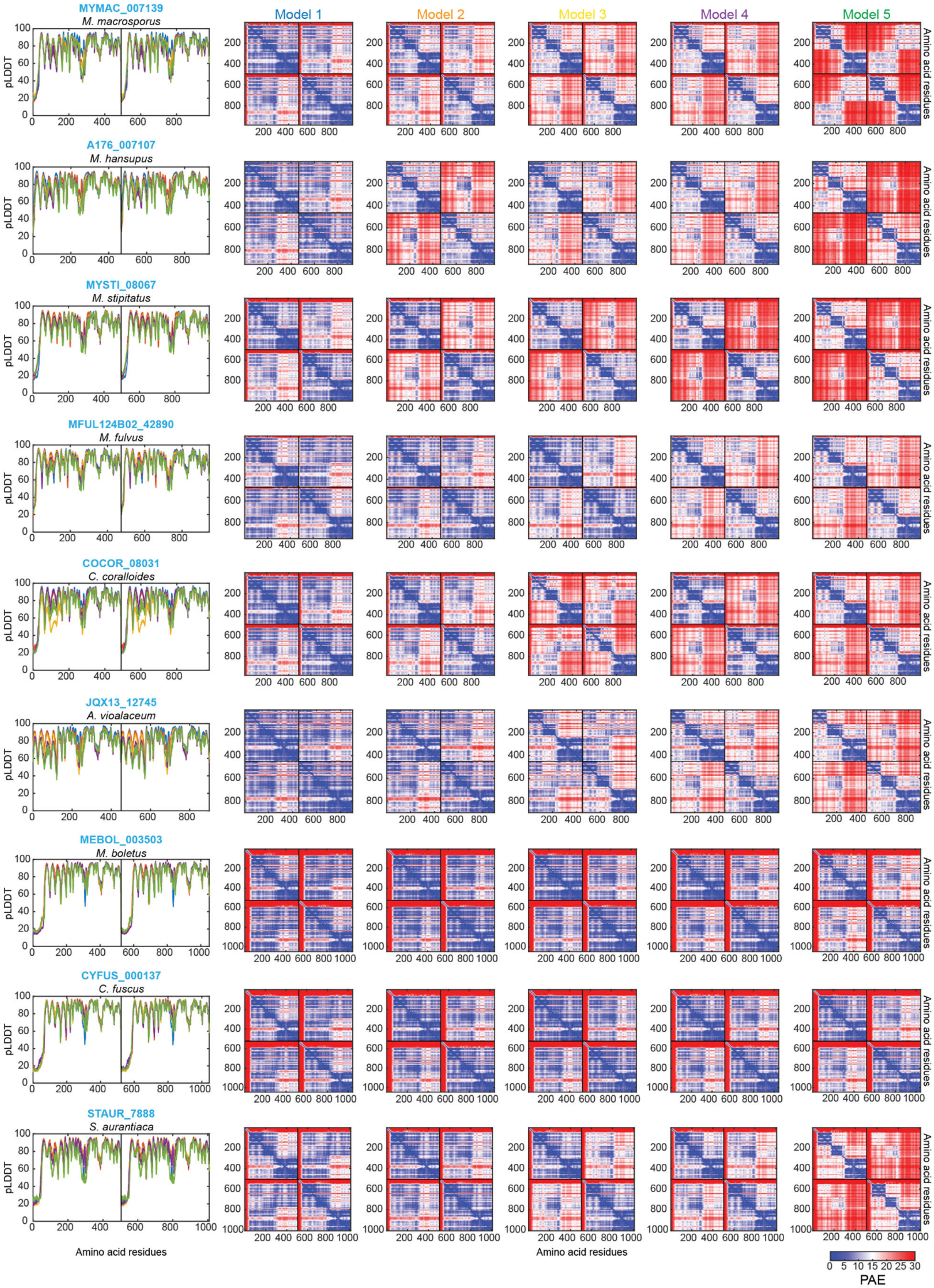

MYMAC_007139
M. macrosporus
A176_007107
M. hansupus
MYSTI_08067
M. stipitatus
MFUL124B02_42890
M. fulvus
COCOR_08031
C. coralloides
JQX13_12745
A. vioalaceum
MEBOL_003503
M. boletus
CYFUS_000137
C. fuscus
STAUR_7888
S. aurantiaca
Model 1
Model 2
Model 3
Model 4
Model 5
pLDDT
Amino acid residues
PAE

Figure S11. AlphaFold2-Multimer models of dimers of nine myxobacterial EpsZ orthologs encoded together with an EpsW ortholog.

Left panels, pLDDT plot shown for the five generated dimeric models of each EpsZ ortholog. Right panels, pAE plots shown for the five generated dimeric models of each EpsZ ortholog. Model rank 1 were selected for further analyses.

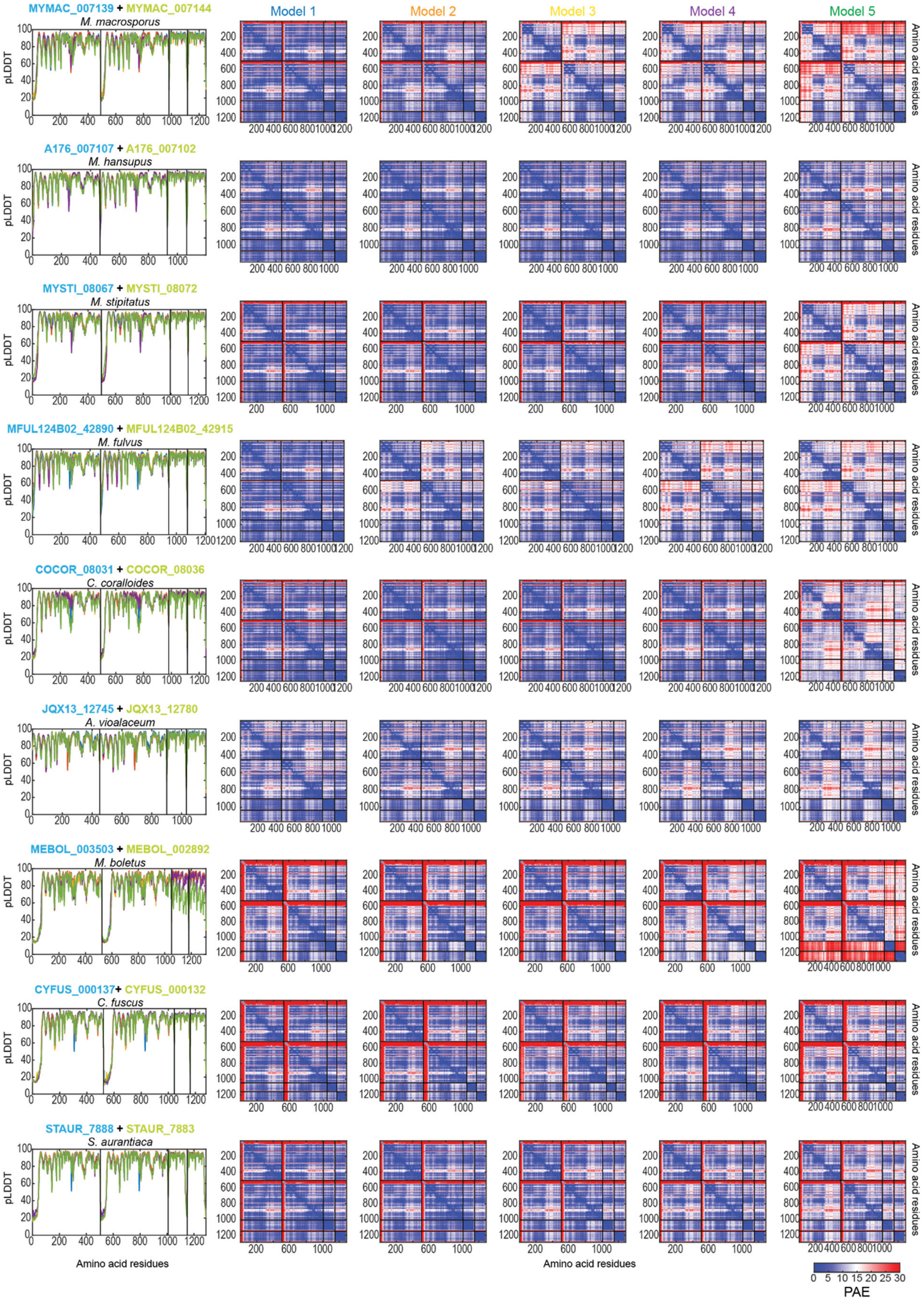
MYMAC_007139 + MYMAC_007144
M. macrosporus
A176_007107 + A176_007102
M. hansupus
MYSTI_08067 + MYSTI_08072
M. stipitatus
MFUL124B02_42890 + MFUL124B02_42915
M. fulvus
COCOR_08031 + COCOR_08036
C. coralloides
JQX13_12745 + JQX13_12780
A. violaceum
MEBOL_003503 + MEBOL_002892
M. boletus
CYFUS_000137+ CYFUS_000132
C. fuscus
STAUR_7888 + STAUR_7883
S. aurantiaca
Model 1
Model 2
Model 3
Model 4
Model 5
pLDDT
Amino acid residues
PAE
0 5 10 15 20 25 30

Figure S12. AlphaFold2-Multimer models of dimers of heterocomplexes of nine myxobacterial EpsZ and EpsW orthologs in 2:2 stoichiometry.

Left panels, pLDDT plot shown for the five generated models of each heterocomplex (2:2 stoichiometry) composed of myxobacterial orthologs of EpsZ and EpsW. Right panels, pAE plots shown for these models. Model rank 1 were selected for further analyses.

**Table S1** Genes with significantly changed transcript levels in the Δ*epsW* mutant compared to WT.

| Gene | Function | $Log_2$ ratio[1] | $-Log_{10}$ *P*-value |
|---|---|---|---|
| *MXAN_0504* | Hypothetical protein | -2.64 | 155.0 |
| *MXAN_1370* | Hypothetical protein | -2.64 | 88.2 |
| *fimU3* | Minor pilin | -2.38 | 71.1 |
| *pilV3* | Minor pilin | -2.37 | 39.0 |
| *pilW3* | Minor pilin | -2.29 | 180.1 |
| *pilX3* | Minor pilin | -2.22 | 56.9 |
| *MXAN_7017* | Csd3, CRISPR-associated | 2.03 | 3.1 |
| *MXAN_7020* | Cas3, CRISPR-associated | 2.13 | 4.1 |
| *MXAN_7018* | Csd1, CRISPR-associated | 2.39 | 3.4 |
| *MXAN_7019* | Cas51, CRISPR-associated | 2.99 | 4.2 |

[1] Numbers indicate $Log_2$ ratio of the mean transcripts in the Δ*epsW* mutant over the mean transcripts in the WT calculated using the DESeq2 method based on four biological replicates per strain. Criteria for significant changes: $Log_2$ (FC) ≥ 2.00 or ≤ -2.00 and $-Log_{10}$ (*P*-value) ≥ 3.0. Note that in this global analysis, we used stringent fold-change cut-offs for quantitative analysis, while in Figure 1C, Figure S1A and B, Figure S2A, and Figure S3A, no fold-change cut-off was used.

**Table S2** Proteins with significantly changed accumulation levels in the Δ*epsW* mutant compared to WT.

| **Protein** | **Function** | **$Log_2$ ratio**[1] | **$-Log_{10}$ *P*-value** |
|---|---|---|---|
| FibA | Matrix-associated zinc metalloprotease FibA | -5.89 | 8.6 |
| MXAN_4534 | Chitinase, class I | -4.14 | 7.1 |
| MsrBA | Methionine-R-sulfoxide reductase/methionine-S-sulfoxide reductase | -3.95 | 3.5 |
| MXAN_1651 | Uncharacterized protein | -3.86 | 6.6 |
| MXAN_3676 | Uncharacterized protein | -3.85 | 9.0 |
| MXAN_6837 | Putative lipoprotein | -3.77 | 4.4 |
| MXAN_0793 | Putative lipoprotein | -3.70 | 8.7 |
| MXAN_2837 | Putative lipoprotein | -3.56 | 4.9 |
| MXAN_7464 | Uncharacterized protein | -3.17 | 4.3 |
| MXAN_1967 | Putative peptidase. S8 (Subtilisin) family | -2.99 | 5.8 |
| MXAN_0225 | Putative long-chain-fatty-acid-CoA ligase | -2.80 | 4.0 |
| MXAN_1672 | Uncharacterized protein | -2.76 | 5.0 |
| MXAN_1287 | Conserved domain protein | -2.74 | 4.4 |
| MXAN_4411 | JmjC domain protein | -2.65 | 3.8 |
| MXAN_0201 | Hydrolase. alpha/beta fold family | -2.63 | 3.4 |
| MXAN_2903 | Putative lipoprotein | -2.59 | 3.8 |
| MXAN_6709 | Putative lipoprotein | -2.35 | 3.8 |
| MXAN_7070 | Uncharacterized protein | -2.34 | 3.5 |
| MXAN_4416 | Cephalosporin hydroxylase family protein | -2.21 | 3.3 |
| MXAN_4524 | Uncharacterized protein | -2.15 | 5.1 |
| MXAN_5308 | Putative lipoprotein | -2.14 | 5.1 |
| MXAN_6116 | Uncharacterized protein | -2.04 | 3.8 |
| MXAN_7039 | Putative lipoprotein | 2.32 | 8.8 |
| MXAN_3114 | DnaJ domain protein | 2.43 | 4.4 |
| MXAN_5560 | Cytochrome c family protein | 2.47 | 3.7 |
| MXAN_2579 | Metallophosphoesterase/PKD domain protein | 2.83 | 3.5 |
| MXAN_5456 | Uncharacterized protein | 3.70 | 4.5 |

[1] $Log_2$ ratio of the mean protein intensities in the Δ*epsW* mutant over the mean protein intensities in the WT calculated based on four biological replicates per strain. Criteria for significant changes: $Log_2$ (FC) ≥ 2.00 or ≤ -2.00 and $-Log_{10}$ (*P*-value) ≥ 3.0. Note that in this global analysis, we used stringent fold-change cut-offs for quantitative analysis, while in Figure 1C, Figure S1A and B, Figure S2A, and Figure S3A, no fold-change cut-off was used.

**Table S3.** Surface accessible, cytoplasmic Lys residues in Eps proteins and DifE[1].

| Protein | Function | Predicted subcellular localization | Surface-accessible, cytoplasmic Lys residues |
|---|---|---|---|
| EpsZ | Large monoPGT | IM | 13 |
| $Wzx_{EPS}$ | Wzx-like flippase | IM | 3 |
| EpsY | $^{D1D2}$OPX | Periplasm | 0 |
| EpsX | Integral OM 18-stranded β-barrel protein | OM | 0 |
| EpsW | Single-domain response regulator | Cytoplasm | 5 |
| EpsV | PCP | IM | 2 |
| EpsU | GT | IM | 12 |
| EpsH | GT | Cytoplasm | 18 |
| $Wzy_{EPS}$ | Wzy-like polymerase | IM | 2 |
| EpsE | GT | Cytoplasm | 13 |
| EcpK | BY pseudokinase | Cytoplasm | 10 |
| EpsD | GT | Cytoplasm | 7 |
| EpsA | GT | Cytoplasm | 11 |
| DifE | Histidine protein kinase | Cytoplasm | 17 |

**Table S4.** Proteins significantly enriched in miniTurbo-EpsW-FLAG proximity labeling experiments in the otherwise WT strain[1].

| Locus Tag | Name | Annotation | $Log_2$ ratio[2] | $-Log_{10}$ *P*-value |
|---|---|---|---|---|
| MXAN_7420 | EpsW | Single-domain response regulator | 11.2 | 6.3 |
| MXAN_0631 | | Transcriptional regulator, AraC family | 7.1 | 5.6 |
| MXAN_2994 | | Glycosyl hydrolase, family 57 | 5.0 | 6.0 |
| MXAN_3056 | | Uncharacterized protein | 4.5 | 4.7 |
| MXAN_6692 | DifE | Histidine protein kinase, Dif system | 4.2 | 4.5 |
| MXAN_6931 | | Thioredoxin | 4.2 | 4.8 |
| MXAN_7415 | EpsZ | Phosphoglycosyl transferase | 4.0 | 4.4 |
| MXAN_4666 | | General secretion pathway protein E, N-terminal domain protein | 3.7 | 6.2 |
| MXAN_5807 | | Putative membrane protein | 3.7 | 4.0 |
| MXAN_4576 | | Acetyltransferase, GNAT family | 3.6 | 5.6 |
| MXAN_2578 | | Methyltransferase, RsmB/NOP family | 3.1 | 4.7 |

[1] EpsW is indicated in green, DifE and EpsZ together with the two other proteins that were enriched in the otherwise WT and the Δ*difD*Δ*difG* strains but not in the Δ*difE* strain are marked in orange and yellow, respectively. Proteins potentially involved in monosaccharide synthesis or modification are marked in blue.

[2] $Log_2$-fold ratios of mean protein intensities in mTurbo-EpsW-FLAG samples relative to the sfGFP-mTurbo-FLAG samples based on four biological replicates. Significantly enriched proteins fulfill the criteria $log_2$ fold ratio ≥3.0; $-log_{10}$ *P*-value ≥4.0.

**Table S5.** Proteins significantly enriched in miniTurbo-EpsW-FLAG proximity labeling experiments in the Δ*difE* background[1].

| Locus Tag | Name | Annotation | $Log_2$ ratio[2] | $-Log_{10}$ *P*-value |
|---|---|---|---|---|
| MXAN_4082 | FusA3 | Elongation factor G3 | 13.2 | 8.8 |
| MXAN_7420 | EpsW | Single-domain response regulator | 12.2 | 4.8 |
| MXAN_3079 | | Phasin family protein | 5.4 | 4.8 |
| MXAN_2993 | | Conserved domain protein | 5.3 | 4.7 |
| MXAN_6798 | | Type I restriction enzyme R protein N-terminal domain-containing protein | 4.6 | 6.0 |
| MXAN_4666 | | General secretion pathway protein E. N-terminal domain protein | 4.2 | 6.5 |
| MXAN_3986 | | ABC transporter, ATP-binding protein | 3.6 | 5.5 |
| MXAN_0731 | | Tryptophan 2.3-dioxygenase | 3.4 | 4.4 |
| MXAN_6931 | | Thioredoxin | 3.2 | 4.1 |
| MXAN_3943 | | Cytochrome P450 family protein | 3.1 | 5.2 |
| MXAN_6765 | | ABC transporter, ATP-binding protein | 3.0 | 8.0 |

[1] EpsW is indicated in green.

[2] $Log_2$-fold ratios of mean protein intensities in mTurbo-EpsW-FLAG samples relative to the sfGFP-mTurbo-FLAG samples based on four biological replicates. Significantly enriched proteins fulfill the criteria $log_2$ fold ratio ≥3.0; $-log_{10}$ *P*-value ≥4.0,

**Table S6.** Proteins significantly enriched in miniTurbo-EpsW-FLAG proximity labeling experiments in the Δ*difD*Δ*difG* background[1].

| Locus Tag | Name | Annotation | $Log_2$ ratio[2] | $-Log_{10}$ *p*-value |
|---|---|---|---|---|
| MXAN_7420 | EpsW | Single-domain response regulator | 14.2 | 7.6 |
| MXAN_4402 | | Non-ribosomal peptide synthetase | 8.1 | 6.0 |
| MXAN_2365 | | Uncharacterized protein | 7.9 | 4.3 |
| MXAN_7298 | | Cytochrome P450 family protein | 7.0 | 4.9 |
| MXAN_0631 | | Transcriptional regulator, AraC family | 7.0 | 7.1 |
| MXAN_3943 | | Cytochrome P450 family protein | 6.8 | 4.4 |
| MXAN_7415 | EpsZ | Phosphoglycosyl transferase | 5.4 | 4.5 |
| MXAN_4292 | | Polyketide synthase | 5.4 | 4.8 |
| MXAN_2949 | | Cation ABC transporter | 5.2 | 5.0 |
| MXAN_2367 | | Acetyltransferase, GNAT family | 5.0 | 5.6 |
| MXAN_6883 | | Dienelactone hydrolase family protein | 4.9 | 4.7 |
| MXAN_3241 | | Uncharacterized protein | 4.8 | 5.3 |
| MXAN_5074 | SpoVG | Putative septation protein SpoVG | 4.4 | 4.1 |
| MXAN_2977 | | Uncharacterized protein | 4.4 | 5.0 |
| MXAN_0282 | | Uncharacterized protein | 4.3 | 4.4 |
| MXAN_4730 | | Lipoprotein releasing system. transmembrane protein, LolC/E family | 4.3 | 4.8 |
| MXAN_2283 | DusA | tRNA-dihydrouridine synthase | 4.2 | 5.7 |
| MXAN_6758 | | Uncharacterized protein | 4.0 | 4.1 |
| MXAN_4410 | | Cephalosporin hydroxylase family protein | 3.9 | 4.0 |
| MXAN_1006 | | Uncharacterized protein | 3.9 | 4.2 |
| MXAN_7398 | | Histidine kinase | 3.8 | 4.3 |
| MXAN_2788 | | Rieske 2Fe-2S domain protein | 3.6 | 6.0 |
| MXAN_2926 | | Ferredoxin. 2Fe-2S | 3.5 | 4.1 |
| MXAN_5807 | | Putative membrane protein | 3.5 | 4.1 |
| MXAN_6692 | DifE | Histidine protein kinase, Dif system | 3.4 | 6.2 |
| MXAN_1308 | | Uncharacterized protein | 3.4 | 8.1 |
| MXAN_4735 | | Putative membrane protein | 3.3 | 5.2 |
| MXAN_3788 | | Uncharacterized protein | 3.3 | 5.5 |
| MXAN_1292 | | Uncharacterized protein | 3.2 | 4.0 |
| MXAN_0429 | | Uncharacterized protein | 3.1 | 5.1 |
| MXAN_5156 | | Uncharacterized protein | 3.1 | 5.5 |
| MXAN_4302 | | FAD-binding domain protein | 3.1 | 4.3 |
| MXAN_1566 | | Uncharacterized protein | 3.1 | 4.5 |

[1] EpsW is indicated in green, DifE and EpsZ together with the two other proteins that were enriched in the otherwise WT and the Δ*difD*Δ*difG* strains but not in the Δ*difE* strain are marked in orange and yellow, respectively. Proteins potentially involved in monosaccharide synthesis or modification are marked in blue.

[2] $Log_2$-fold ratios of mean protein intensities in mTurbo-EpsW-FLAG samples relative to the sfGFP-mTurbo-FLAG samples based on four biological replicates. Significantly enriched proteins fulfill the criteria $\log_2$ fold ratio ≥3.0; $-\log_{10}$ *P*-value ≥4.0,

**Table S7.** Strains used in this work

| Species and strain | Genotype | Reference or source |
|---|---|---|
| *M. xanthus* | | |
| DK1622 | WT | (48) |
| DK10410 | Δ*pilA* | (42) |
| SA7400 | Δ*epsZ* | (49) |
| SA6888 | Δ*cluster*_1Δ*cluster*_2 | (50) |
| SA6892 | Δ*cluster*_2Δ*cluster*_3 | (50) |
| SA6899 | Δ*cluster*_1Δ*cluster*_3 | (50) |
| SA7609 | Δ*cluster*_1Δ*cluster*_2Δ*cluster*_3 | (50) |
| SA5649 | Δ*difE* | This work |
| SA7415 | Δ*epsW* | This work |
| SA8550 | Δ*epsW attB*::pMP146 *(*$P_{pilA}$ *epsW)* | This work |
| SA11564 | Δ*difD*Δ*difG* | This work |
| SA11578 | Δ*difD*Δ*difG*Δ*epsW* | This work |
| SA11639 | Δ*epsW attB*::pJSc105 *(*$P_{pilA}$ *mTurbo-epsW-FLAG)* | This work |
| SA11646 | Δ*epsW attB*::pJSc113 *(*$P_{pilA}$ *epsW) 18-19*::pMH97 *(*$P_{van}$ *sfGFP-mTurbo-FLAG*) | This work |
| SA7425 | Δ*difE*Δ*epsW* | This work |
| SA11645 | Δ*difE*Δ*epsW attB*::pJSc105 *(*$P_{pilA}$ *mTurbo-epsW-FLAG))* | This work |
| SA11649 | Δ*difE*Δ*epsW attB*::pJSc113 *(*$P_{pilA}$ *epsW)* | This work |
| SA11647 | Δ*difE*Δ*epsW attB*::pJSc113 *(*$P_{pilA}$ *epsW) 18-19*::pMH97 *(*$P_{van}$ *sfGFP-mTurbo-FLAG*) | This work |
| SA11644 | Δ*difD*Δ*difG*Δ*epsW attB*::pJSc105 *(*$P_{pilA}$ *mTurbo-epsW-FLAG)* | This work |
| SA13200 | Δ*difD*Δ*difG*Δ*epsW attB*::pJSc113 *(*$P_{pilA}$ *epsW)* | This work |
| SA11648 | Δ*difD*Δ*difG*Δ*epsW attB*::pJSc113 *(*$P_{pilA}$ *epsW) 18-19*::pMH97 *(*$P_{van}$ *sfGFP-mTurbo-FLAG*) | This work |
| SA13207 | Δ*epsW attB*::pJSc146 *(*$P_{pilA}$ *epsW-His*$_{10}$*)* | This work |
| *Salmonella* | | |
| LT2 | *S. enterica* serovar Typhimurium, WT | S. Maloy |
| MSS2 | LT2, Δ*wbaP*::*cat* | (51) |
| JMF20 | LT2, Δ*wbaP* | This work |
| *E. coli* | | |
| *E. coli* NEB Turbo | F' *proA*$^{+}$*B*$^{+}$ *lacI*$^{q}$ *ΔlacZM15 / fhuA2 Δ(lac-proAB) glnV galK16 galE15 R(zgb-210::Tn10)*Tet$^{S}$ *endA1 thi-1 Δ(hsdS-mcrB)5* | New England Biolabs |

**Table S8.** Plasmids used in this work

| Plasmid | Description | Reference or source |
|---|---|---|
| pBJ114 | *galK* $Kan^R$ | (52) |
| pSW105 | $Km^r$ $P_{pilA}$ | (38) |
| pSWU30 | $Tet^r$ | (42) |
| pBAD33 | Arabinose-inducible promoter, $Cam^r$ | (53) |
| pBADNTF | pBAD24 for N-terminal FLAG fusion and with arabinose-inducible promoter, $Amp^r$ | (54) |
| pMP036 | pBJ114, in-frame deletion construct for *epsW* $Km^r$ | This work |
| pDJS102 | pBJ114, in-frame deletion construct for *difE* $Km^r$ | This work |
| pJSc002 | pBJ114, in-frame deletion construct for *difD* $Km^r$ | This work |
| pJSc003 | pBJ114, in-frame deletion construct for *difG* $Km^r$ | This work |
| pJSc105 | pSW105, $P_{pilA}$ *mTurbo-epsW-FLAG* $Km^r$ | This work |
| pJSc113 | pSWU30, $P_{pilA}$ *epsW* $Tet^r$ | This work |
| pJSc143 | pBAD33, *epsW-His$_{10}$*, $Cam^r$ | This work |
| pJSc146 | pSW105, $P_{pilA}$ *epsW-His$_{10}$* | This work |
| pMP145 | pSW105, $P_{pilA}$ *epsW* $Km^r$ | This work |
| pMP146 | pBADNTF, *FLAG-epsZ* $Amp^r$ | (49) |
| pMH97 | pMR3690, $P_{Van}$ *sfGFP-mTurbo-FLAG* $Km^r$ | (1) |
| pSM13 | pUC18, *wbaP* from *S. enterica* containing a 1-bp deletion at position 583 and a 2-bp deletion at position 645, which causes a frame shift at WbaP I194 and frame restoration at Y215, $Amp^r$ | (51) |
| pCP20 | FLP recombinase expression, $Cam^r$, $Amp^r$ | (55) |

**Table S9.** Oligonucleotides used in this work[1]

| Primer name | Sequence 5'-3' | Brief description |
|---|---|---|
| 7420_A | ATCGGGTACCAGTCGTGGGGACAGCATCCT | For Δ*epsW* |
| 7420_B | CCCAGTGCTGAGGACCGTGTGCAGGTC | For Δ*epsW* |
| 7420_C | ACGGTCCTCAGCACTGGGGCCAGGAGG | For Δ*epsW* |
| 7420_D | ATCGTCTAGACAGTACTCCACCAGCTTCGC | For Δ*epsW* |
| 7420_E | ACCTGACGCTGACGGGGACC | For Δ*epsW* |
| 7420_F | CACCTTCTCCTTCAGCCGGT | For Δ*epsW* |
| 7420_G | GGTGGAGGACGCCCCCTTCT | For Δ*epsW* |
| 7420_H | TGAGGAGCGACTGCACCCGC | For Δ*epsW* |
| 7420_$P_{pilA}$ fw | ATCGTCTAGAATGGAGTCCGACCTGCACAC | For complementation of *epsW* |
| 7420_$P_{pilA}$ rev | ATCGAAGCTTTCACGGTGCCCTCCTGGCCC | For complementation of *epsW* |
| mTurbo-$P_{pilA}$-for | GCGCTCTAGAATGATCCCGCTCCTG | For mTurbo-EpsW-FLAG |
| mTurbo-epsW-FLAG int for | GGCGGGGGCGGGAGCGAGTCCGACCTGCAC<br>ACGG | For mTurbo-EpsW-FLAG |
| mTurbo-epsW-FLAG int rev | GCTCCCGCCCCCGCCCTTCTCGGCGCTGCGCA | For mTurbo-EpsW-FLAG |
| EpsW-FLAG-rev | CAGTAAGCTTTCACTTGTCGTCGTCGTCCTTGTA<br>GTCCGGTGCCCTCCTGGCCCCAG | For mTurbo-EpsW-FLAG |
| EpsW-His-rev-HindIII | ACTGAAGCTTTCAGTGGTGGTGGTGATGGTGATG<br>ATGATGATGCGGTGCCCTCCTGGCCCC | For EpsW-$His_{10}$ |
| 6692_A | ATCGGAATTCAGGCTGGGTGGCCTGGGAGT | For Δ*difE* |
| 6692_B | CGGCACGTCGAGGTAGCGGGACATGTC | For Δ*difE* |
| 6692_C | CGCTACCTCGACGTGCCGAGGTTACTG | For Δ*difE* |
| 6692_D | ATCGTCTAGACAGCAGGTCGCGGATGCTGT | For Δ*difE* |
| 6692_E | ACGGCGCCAGGAAAGATGAC | For Δ*difE* |
| 6692_F | TTTCAGCGCCTTCTCATCCA | For Δ*difE* |
| 6692_G | AAGGTGACGGCGCTGAAGCC | For Δ*difE* |
| 6692_H | CCAGCGCTGGGCCAGCGTTT | For Δ*difE* |
| 6693_A | AAAGGTACCTGCAAGGACGTCGGCGGC | For Δ*difD* |
| 6693_B | CTTCTTCACGTCGACGACCAGGACCCG | For Δ*difD* |
| 6693_C | GTCGTCGACGTGAAGAAGGTCCTGGGC | For Δ*difD* |
| 6693_D | TTTTCTAGAAGCTTCCACCAGCTCCGC | For Δ*difD* |
| 6693_E | GCAGAAGGTGCTCAAGGC | For Δ*difD* |
| 6693_F | AGGGAGTTGGCCAGGTCG | For Δ*difD* |
| 6693_G | GGACATCTTTGCGTCTGG | For Δ*difD* |
| 6693_H | GACGATGAAGTCCGAGGC | For Δ*difD* |
| 6691_A | TTTGGTACCGGAGGCGCCCGAGGATCG | For Δ*difG* |
| 6691_B | CAGCAGGTCCGCGTCGCTGTGCAGGGG | For Δ*difG* |
| 6691_C | AGCGACGCGGACCTGCTGGCGCGGCTG | For Δ*difG* |

| | | |
|---|---|---|
| 6691_D | TTT<u>TCTAGA</u>GCATGGTGCTCACCTGCG | For Δ*difG* |
| 6691_E | ACACGTTGGTGAAGGACC | For Δ*difG* |
| 6691_F | ATGTCGTCGATGGACACG | For Δ*difG* |
| 6691_G | GCAAGGTGGACCTCTCCA | For Δ*difG* |
| 6691_H | TCCAACACCACCACGCTG | For Δ*difG* |

[1] Underlined sequences indicate restriction sites.

**Table S10.** Fully sequenced myxobacterial genomes used for the 16S RNA tree and gene co-occurrence analysis.

| Species and strain name |
|---|
| *Anaeromyxobacter dehalogenans* 2CP-C |
| *Anaeromyxobacter* sp. Fw109-5 |
| *Anaeromyxobacter* sp. K |
| *Archangium gephyra* DSM 2261 |
| *Archangium violaceum* Cb SDU34 |
| *Chondromyces crocatus* Cm c5 |
| *Corallococcus coralloides* DSM 2259 |
| *Cystobacter fuscus* DSM 52655 |
| *Haliangium ochraceum* DSM 14365 |
| *Labilithrix luteola* DSM 27648 |
| *Melittangium boletus* DSM 14713SG |
| *Minicystis rosea* DSM 24000 |
| *Myxococcus fulvus* 124B02 |
| *Myxococcus macrosporus* DSM 14675 |
| *Myxococcus hansupus* (*Myxococcus* sp. mixupus) |
| *Myxococcus stipitatus* DSM 14675 |
| *Myxococcus xanthus* DK1622 |
| *Sandaracinus amylolyticus* DSM 53668 |
| *Sorangium cellulosum* So ce 56 |
| *Stigmatella aurantiaca* DW4/3-1 |
| *Vulgatibacter incomptus* DSM 27710 |
